# Pyruvate kinase allostery resist hyperglycemia, obesity and inflammation by inducing TCA vortex and glucose U turn

**DOI:** 10.1101/2024.08.09.607407

**Authors:** Xuan Zhang, Xudong Jiang, Xiaobin Wu, Youhao Yang, Jintian Wang, Junfeng Zheng, Miao Zhou, Qian Chen

## Abstract

ATP is the final product of glycolysis and the TCA cycle. However, the counter-regulation of glycosis and TCA by ATP is poorly understood. Here we show that ATP analog celastrol (CLT) binds to the ATP binding pocket on pyruvate kinase PKM (PKM) while inducing allosteric regulation of PKM. Quantum calculation predicts hydrogen bond formation between CLT and asparagine. Liquid chromatography-mass spectrometry further identifies PKM as CLT’s target. The inhibition of PKM is stronger and longer for CLT while weak and short for ATP. Notably, CLT-PKM interaction perfectly underlies the hypoglycemic effects by forming glucose carbon flux U turn before PKM. Besides, the PKM allostery induces a tricarboxylic acid (TCA) vortex which could promote amino acid and lipid degradation as the energy compensation, leading to a significant weight loss. Additionally, CLT exerts efficient antioxidant effects by altering the glucose flux to strengthen the pentose phosphate pathway. Consequently, the CLT-PKM interaction vividly reproduces the ATP-PKM interaction named “ATP resistance” in the diabetes pathogenesis.

## Introduction

Plants and animals have evolved a wonderful complementary relationship in supplying and consuming carbon such as glucose as the major energy source. Main metabolites in the central carbon metabolism tricarboxylic acid (TCA) cycle have plant root. Citric acid in the citrus fruits (Huang et al., 2023), succinic acid in the resinite (Poulin and Helwig, 2014), fumaric acid in the corydalis (Coffey and Simon, 2024) and malate in the apple (Gao et al., 2024) have shown various and amazing physiological and medical implications. Meanwhile, plant extracts and herbs continue to provide front-line pharmacotherapy for many millions of people worldwide (Corson and Crews, 2007). Successful example such as metformin derived from Galega officinalis has been the first-line medication to treat type 2 diabetes mellitus and is used daily by >200 million patients (Foretz et al., 2023). However, direct targets and underlying molecular mechanism of many other promising candidates like celastol (CLT) still remain elusive (Corson and Crews, 2007). Just like cellular energy charge alteration glucoregulatory effect by metformin, CLT could also protect against obesity and metabolic dysfunction (Liu et al., 2015; Ma et al., 2015). Surprisingly, the mechanisms underlying how they interact with glucose and other energy metabolism are still not fully understood (Foretz et al., 2019).

Metabolic disorder and excessive glucose uptake are the main reasons for chronic diseases such as diabetes and obesity with unclear cellular mechanisms (Klein et al., 2022). Hopefully, increasing attention has been focused on important evidence including the fact that glycolysis has high control strength over insulin secretion and pyruvate kinase has favorable bioenergetics for raising ATP (Merrins et al., 2024). In addition, pyruvate kinase PKM (PKM) is the rate-limiting enzyme to control glycolysis and central to glucose metabolism and energy expenditure (Chen et al., 2019). In this process, ATP is produced by phosphoenolpyruvate (PEP) and ADP substrate level phosphorylation (Abulizi et al., 2020). However, excessive ATP could in turn inhibit PKM (Callens et al., 1991). Rate-limiting enzymes in the glycolytic pathway including PKM are currently attracting increasing attention in regulating blood glucose level (Corkey, 2020; Lewandowski et al., 2020; Starling, 2021). In addition, PKM could regulate reactive oxygen species (ROS) (Toller-Kawahisa et al., 2023) and play an important role in inflammation. However, several important questions still remain unclarified. Does cellular glucose flux contribute to the high blood glucose in diabetes other than insulin resistance? Why urine glucose comes into existence? What’s the molecular mechanism of the body weight loss for diabetics? More work needs to be done on glycolysis as well as PKM to understand diabetes better.

Nevertheless, a wide gap exists between the candidate drugs and their potential targets. It is extremely challenging to accurately predict how candidate drugs interact with their targets involving thousands of atoms (Santagati et al., 2024). Hopefully, quantum calculation could analyze the chemical and physical interactions between molecules and atoms, including compounds with amino acids (Ajagekar and You, 2023; Ramesh et al., 2024; Wasielewski et al., 2020). In this study, quantum chemistry and physics combined with other technologies, such as high-performance, high-throughput liquid chromatography-mass spectrometry (LC-MS) proteomics, isotope labeled targeted plus untargeted metabolomics, single-cell RNA-sequencing (scRNA-seq), molecular docking etc., remarkably identify PKM as CLT’s target and reproduce the interaction mechanism between ATP and PKM.

## Results

### 1. CLT shows the strongest interaction with asparagine (Asn) and Serine (Ser) among 20 Amino acids

To theoretically explore the interaction between CLT and potential protein targets, the interactions between CLT and 20 common amino acids were first simulated by density functional theory (DFT) calculations (Bosoni et al., 2024; Jain et al., 2016). Figure 1A shows the scheme of calculation process. To identify the characteristic patterns of positive and negative potentials, the molecular electrostatic potential (MEP) of CLT and 20 common amino acids was calculated by Gaussian as described previously (Cruz et al., 2023; Suresh et al., 2022). Figures 1B, C&D show the MEP of CLT, Asn and Ser as examples respectively. The red represents the electronegative density region, while the electropositive charge is distributed in blue region. The charge density redistribution and density of states (DOS) in Figures 1E-H. Figures S1 & S2 are shown to elucidate their interaction properties (Chen et al., 2020; Jain et al., 2016). Among all interactions, we found that the systems containing Ser or Asn exhibit the strongest binding. The calculated charge density redistribution for interaction between Ser and CLT is shown in Figure 1E. The green denotes electron accumulation, and the yellow color represents depletion. The distance between upper pair oxygen atom and hydrogen atom is 1.73 Å, and the distance for lower pair is 1.624 Å, which strongly indicate the formation of hydrogen bonds. The adsorption energy for this system is - 0.508 eV. As shown in Figures 1F and 1G, results on DOS suggest the hybridization of these two pairs of oxygen-hydrogen bond, as indicated by the overlap of grey and green curve close to the Fermi level. Similar behaviors were also found in the system of CLT and Asn (Figure 1H). The distance for the upper and lower pair oxygen atom and hydrogen atom are 1.824 Å and 1.786 Å, respectively. The adsorption energy is - 0.442 eV. Moreover, electron depletion and accumulation are depicted. The hybridization of these two pairs of oxygen-hydrogen bond is shown in Figures 1I and J. The interactions of CLT with other 18 amino acids are shown in Figures S1 and S2. Strong interaction is also found in the systems containing glutamine (Gln), glutamate (Glu) and tyrosine (Tyr), with adsorption energy larger than -0.4 eV. While the interactions of CLT with alanine (Ala), cysteine (Cys), isoleucine (Ile), leucine (Leu), lysine (Lys), methionine (Met), phenylalanine (Phe), proline (Pro) and valine (Val) are relatively weak in the present reaction position, with adsorption energy smaller than - 0.2 eV. Therefore, the theoretical results show that CLT tends to strongly interact with Ser, Asn and Tyr. These amino acids with strong interactions with CLT might be the possible interacting site in vivo.

**Figure 1.**
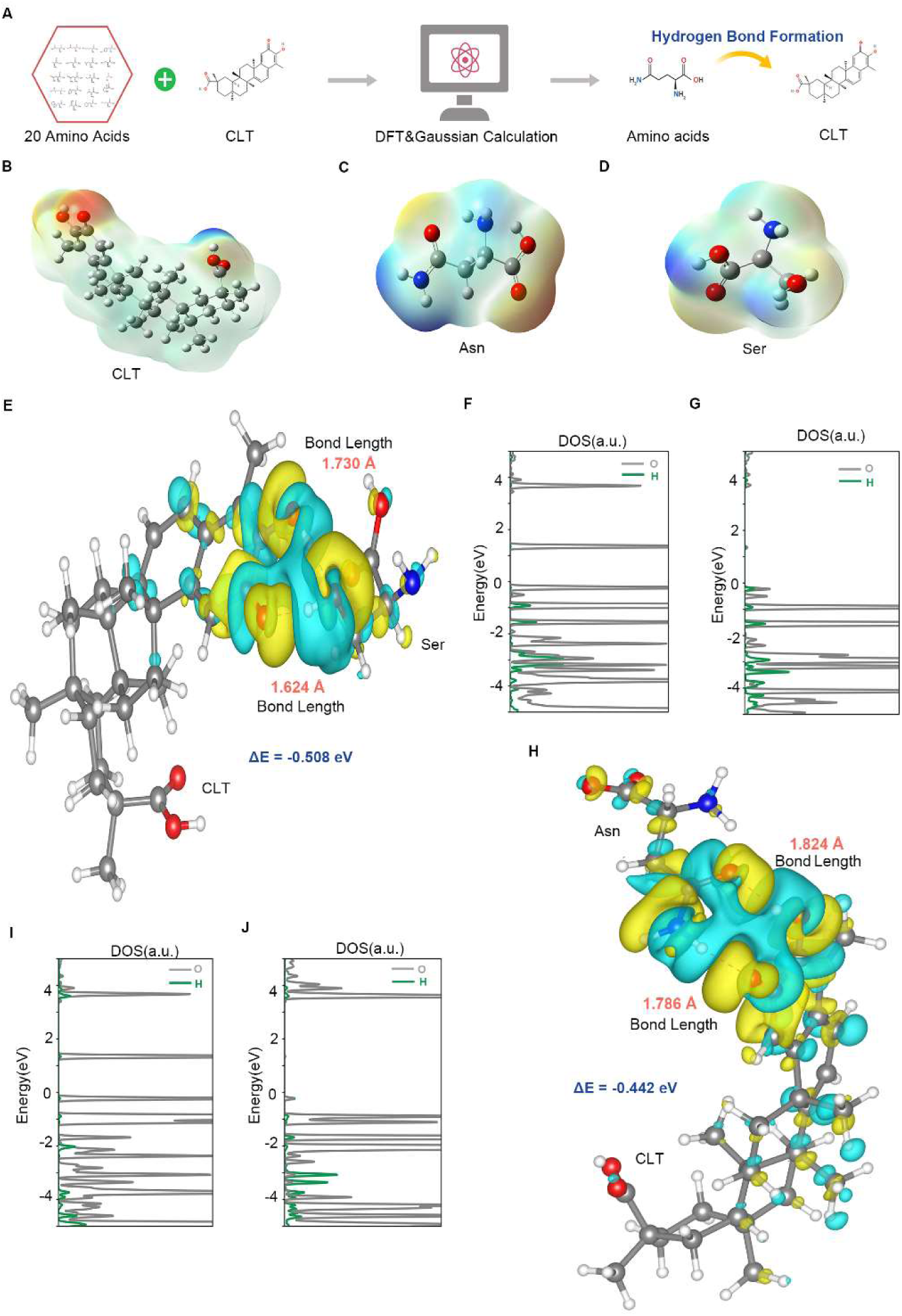
Strong interactions between CLT and Asn, Ser as recognized by quantum calculation. (A) Diagram of computational methods used to simulate interactions between CLT and 20 amino acids. (B) The MEP surface map of CLT. (C&D) The MEP surface map of Asn (C) and Ser (D). (E) Calculated charge density redistribution for interaction of Ser and CLT. (F&G) Projected density of states of selected atom for CLT with Ser. (H) Calculated charge density redistribution for interaction of Asn and CLT. (I&J) Projected density of states of selected atom for CLT with Asn.

### 2. CLT binds to the ATP acting pocket in PKM

To identify the real binding site of CLT in vivo, the proteomics analysis using liquid chromatography-mass spectrometry (LC-MS) was conducted on tissues treated with CLT. The results show that CLT binds to the peptide fragment of pyruvate kinase PKM (PKM). Figures 2A and 2B demonstrate the schema and sequence of amino acids, where green indicates peptide sequence recognized by proteomics. Specifically, CLT was identified to bind to amino acids Asn210 and Pro212 in PKM (as marked by a red C above the selected amino acid). Consistently, the LC-MS binding results are highly consistent with the quantum simulation of CLT interaction with Asn (Figure 1H), encouraging us to further elucidate how CLT interacts with PKM.

**Figure 2.**
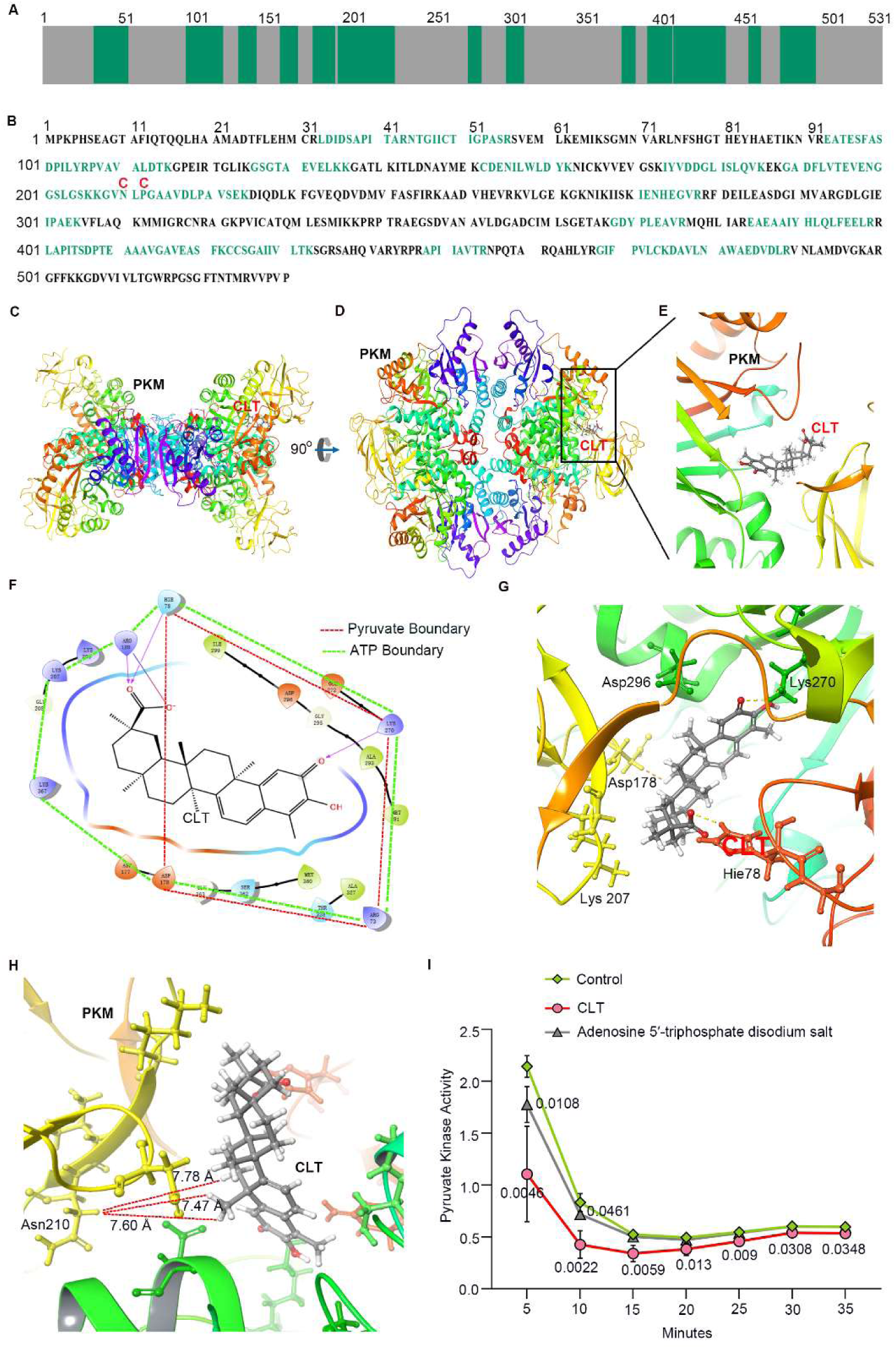
CLT binds to the ATP acting pocket of PKM and inhibits its enzyme activity just like ATP. (A) Scheme of peptide sequence recognized by LC-MS. (B) Amino acid sequence of PKM. CLT modification of PKM was identified at Asn210 and Pro212 as marked red C above the selected amino acid. (C, D&E) Position of CLT in the pocket on PKM. (F) The binding pocket of CLT compared to pyruvate and ATP. (G) The interaction of CLT with amino acids in the binding pocket. (H) The position relationship between CLT and Asn210 and their shortest distance in the binding pocket. (I) CLT shows stronger and longer inhibition of pyruvate kinase activity in comparison to adenosine 5′-triphosphate disodium salt. n=4, the p value compared to control group for each point is listed aside.

Next, protein-ligand interaction of CLT and PKM was simulated in Maestro (Li et al., 2018). The docking score for the CLT-PKM complex is -3.761. Figures 2C and 2D show the side and top view of the PKM-CLT complex. The enlarged view of CLT binding site is shown in Figure 2E. Intriguingly, CLT’s binding pocket is identical to that of ATP (PDB: 4FXF,) and pyruvate (PDB:4YJ5) (Chen et al., 2019) as shown in Figures 2F and 2G. Typically, collision cross sections (CCSs) from ion mobility mass spectrometry measurements are routinely adopted to compare the protein structures and compounds (Landreh et al., 2020). In fact, the CCSs for [CLT+H]+ vary from 202.39 Å² to 205.6 Å² , and the CCSs for [ATP+H]+ range from 196.4 Å² to 196.6 Å². Respectively, the CCSs for [ADP+H]+ and [PEP+Na]+ are 184.8 Å² and 138.1 Å² (Figure S3A) (Picache et al., 2019). Definitely, CLT and ATP are much of a muchness concerning their CCSs. Notably, the PKM binding pocket for CLT (Figure S3B) is exactly the pocket where ATP and pyruvate generate from PEP and ADP (Starling, 2021). Accordingly, the catalytic activity of rate-limiting enzyme PKM should be interrupted by the binding of CLT and ATP (Figure 2I). Generally, CLT binds to the ATP acting pocket of PKM (Figures S3B and S3C) and shows stronger and longer inhibition of PKM activity in comparison to ATP.

Figure 2H shows the position relationship between CLT and Asn210 within the binding pocket on PKM. The shortest distance between CLT and Asn210 (7.47 Å) provides a window of opportunity for CLT to interact with Asn210 (Figure 1H) when the proteins are fragmented into peptides during the preparation step for proteomics (Figures 2A and 2B). The charge surface analysis of PKM-CLT binding complex is shown in Figures S3D and S3E. The blue indicates negative-charged and the red for positive-charged on PKM, while the green indicates negative-charged and the red for positive-charged on CLT. Figure S3E shows that the binding pocket is mainly negative-charged (blue). Meanwhile, liposoluble CLT has poor water-solubility (Shi et al., 2020) and therefore mainly negative-charged (Figures S3D and S3E). In addition, the inside view of the binding pocket in Figure S3F shows it is surrounded by Ile51, Ile88, Leu74, Leu112, glycine79 (Gly79), histidine78 (His78), Ser362 and arginine73 (Arg73). Notably, Ile, Leu, Gly, Ser and Arg are all aliphatic amino acids, constituting the negative-charged binding pocket. Therefore, the liposoluble CLT could easily bind to the ATP-acting pocket surrounded by aliphatic amino acids, as a stiff “ATP” with an appropriate size.

### 3. CLT inhibits glycolysis while decreasing the blood glucose

Since PKM is one of the rate-limiting enzymes for cell glycolysis (Toller-Kawahisa et al., 2023), we next examined the impact of CLT-PKM interaction on glycolysis in A549 cells. Figures 3A-D show that CLT could decrease the glycolytic capacity and the glycolytic reserve in extracellular acidification rate (ECAR), while the knockout of PKM has a more significant inhibitive effect. For oxygen consumption rate (OCR), CLT could inhibit both maximal respiration and spare capacity. Besides, PKM knockout could significantly inhibit basal respiration as well. For CLT, PKM-/- and PKM-/- + CLT, all the major ECAR and OCR index show a steady decline. However, ATP-linked respiration showed no significant changes upon these treatments. Western blotting of PKM protein in different A549 cell groups is shown in Figure 3E. These data prove that CLT-PKM interaction could inhibit the cellular glycolysis.

**Figure 3.**
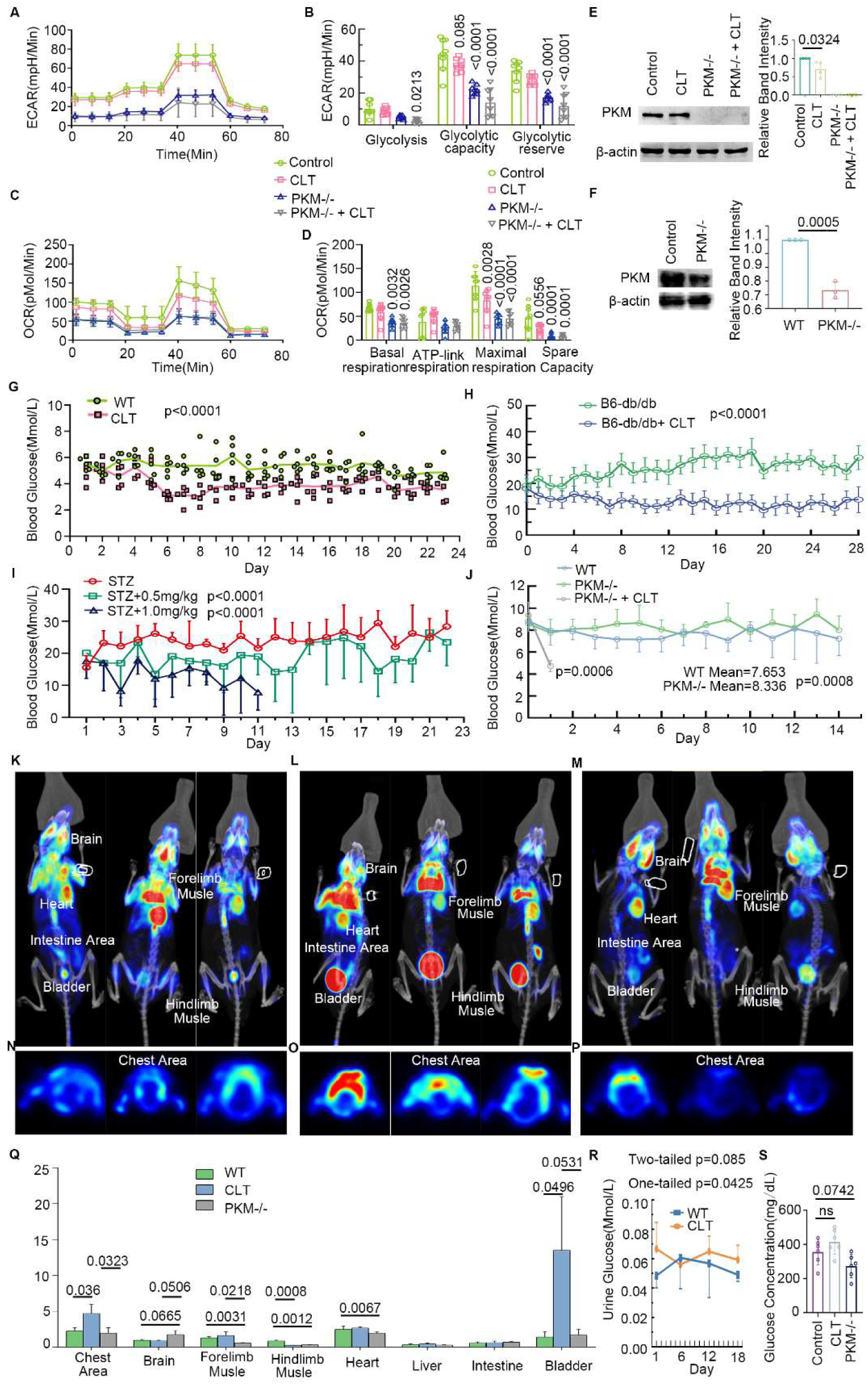
CLT inhibits glycolysis while reducing blood glucose levels. (A-D) Inhibition of ECAR and OCR by CLT and PKM -/- in A549 cells. n=8, each p value is listed in the chart. (E) Western blotting of PKM protein in Control, CLT, PKM-/- and PKM-/- + CLT A549 cell. n=3, p value is listed in the chart. (F) Western blotting of PKM protein in WT and PKM-/- mice. n=3, p=0.0005. (G) CLT reduces the blood glucose levels in WT mice. n=3, p<0.0001. (H) CLT lowers the blood glucose levels in B6-db/db mice. n=12 (B6-db/db), n=16 (B6-db/db + CLT), p<0.0001. (I) CLT downregulates blood glucose levels in STZ mice. n=4 (STZ), n=5 (STZ + 0.5 mg/kg CLT), n=9 (STZ + 1 mg/kg CLT), p<0.0001. (J) PKM knockout mice show a slight increase in blood glucose level and one-day CLT treatment decreases the blood glucose levels effectively. n=6 (WT), n=7 (PKM-/-), n=5 (PKM-/- + CLT), p=0.0008 for PKM-/- group in comparison to WT group, p=0.0006 for PKM-/- + CLT group in comparison to PKM-/- group on Day 1. (K-M) Pet/CT imaging of 18F-FDG distribution in the brain, muscle, heart, liver, intestine, bladder for WT, CLT and PKM-/- mice. (N-P) Pet/CT imaging of 18F-FDG distribution in the chest area for WT, CLT and PKM mice. (Q) Data analysis of 18F-FDG Pet/CT imaging in I-N. (R) Urine glucose levels of WT mice and CLT-treated WT mice. n=5 (WT), n=6 (CLT), p=0.085 for two-tailed t-test and p=0.0425 for one-tailed t-test. (S) In A549 cells, 48h CLT treatment (short PKM inhibition) tends to increase the glucose levels while PKM knockout (strong and long PKM inhibition) is inclined to reduce the glucose levels in the cell culture medium. n=6, each p value is listed in the chart.

Hence, we expected to see the glucose upregulation in the mouse blood. Therefore, wild type (WT) C57 BL/6J mice were treated with CLT via tail vein injection for 3 weeks and the blood glucose levels were monitored daily. Surprisingly, the blood glucose levels went down steadily from the sixth day (Figure 3G). Importantly, CLT could also downregulate blood glucose level in C57BL/6J B6-db/db mice (Figure 3H). Furthermore, CLT could also reduce the blood glucose levels in streptozotocin (STZ) induced type 1 diabetes mouse model (Figure 3I), implicating that its impact on blood glucose is independent of pancreatic islet or insulin. However, PKM-/- mice (PKM protein expression in Figure 3F) showed a slight increase in the blood glucose levels. Subsequently, one-day CLT injection lowered the blood glucose levels immediately (Figure 3J). To this end, the inhibition of glycolysis by PKM and the decrease in blood glucose look unexpectedly contradictory and need further exploration.

Therefore, mice injected with 18F-flurodeoxyglucose (18F-FDG) were assessed on PET/CT scanner (He et al., 2022; He et al., 2019) to demonstrate the glucose uptake and distribution in WT mice, CLT-treated mice and PKM-/- mice (Figures 3K-Q). In CLT-treated mice, chest area rich in brown fat showed a significant increase in 18F-FDG signal (Figure 3O). Notably, a super high signal was found in the bladder in CLT-treated mice (Figure 3L), indicating a surge of urine glucose. Indeed, the increase of urine glucose following CLT treatment was further confirmed by urine glucose assay (Figure 3R). However, the glucose uptake has different patterns in different tissue. The significant higher 18F-FDG signal was only found in the brain region of PKM-/- mice. For forelimb and hindlimb muscles, there was a significant 18F-FDG signal reduction in PKM knockout mice compared to CLT or WT group (Figures 3K-M). The bladder signal of 18F-FDG showed a slight increase without significance when comparing PKM-/- group to the WT group (Figures 3K and 3M). The reason for glucose uptake pattern difference would be analyzed and discussed in the discussion part.

Proteomics analysis was carried out to access the protein changes in CLT-treated WT mice. Remarkably, the string analysis of 1023 proteins surrounded PKM as the network centroid (Figure 4A), further indicating that PKM is the target of CLT. Additionally, the heatmap showed significant different patterns in WT and WT+CLT mouse kidney as shown in Figure 4B. Meanwhile, the string analysis of 713 significantly altered showed similar PKM-centered protein-protein interaction (PPI) network in CLT-treated db/db mice (Figures 4C and 4D). In addition, metabolomics and ^13^C-labeled glucose metabolomics results showed significant changes in metabolite levels in Control, CLT, PKM-/- and PKM-/- + CLT A549 cells. Proteomics and metabolomics analyses (Figures 4E and 4F) are expected to explain the puzzled glucose transporting dynamics. As shown in Figure 5A, the downregulation of glucose levels in CLT, PKM-/- and PKM-/- + CLT group was accessed by ^13^C labeled glucose metabolic flux. In contrast, the ^13^C exchange rates of glucose all increased in CLT, PKM-/- and PKM-/- + CLT group. Moreover, the level and exchange rate of fructose-6-phosphate in Figure 5B were both increased in CLT, PKM-/- and PKM-/- + CLT group. And the increased uptake of 2-deoxy-2-[7-nitro-2,1,3-benzoxadiazol-4-yl)amino]-D-glucose (2-NBDG) in HK-2 cells upon CLT treatment was also validated by flow cytometry (Figure 5C). Taking together, these results indicate the increase of glucose uptake by CLT and PKM knockout. Nevertheless, the flow direction of increased uptake of glucose needs further elucidation. Since glycolysis as well as PKM is vital in cell energy metabolism (Wang et al., 2023; Wei et al., 2023), the ATP level was therefore reduced by CLT treatment (Figure 5D). Meanwhile, the key ATP resynthesis source phosphocreatine (Hargreaves and Spriet, 2020) was also decreased in PKM knockout and CLT treated PKM knockout cells (Figure 5E). Taking together, the increased uptake of glucose might be the cellular response to the energy shortage.

**Figure 4.**
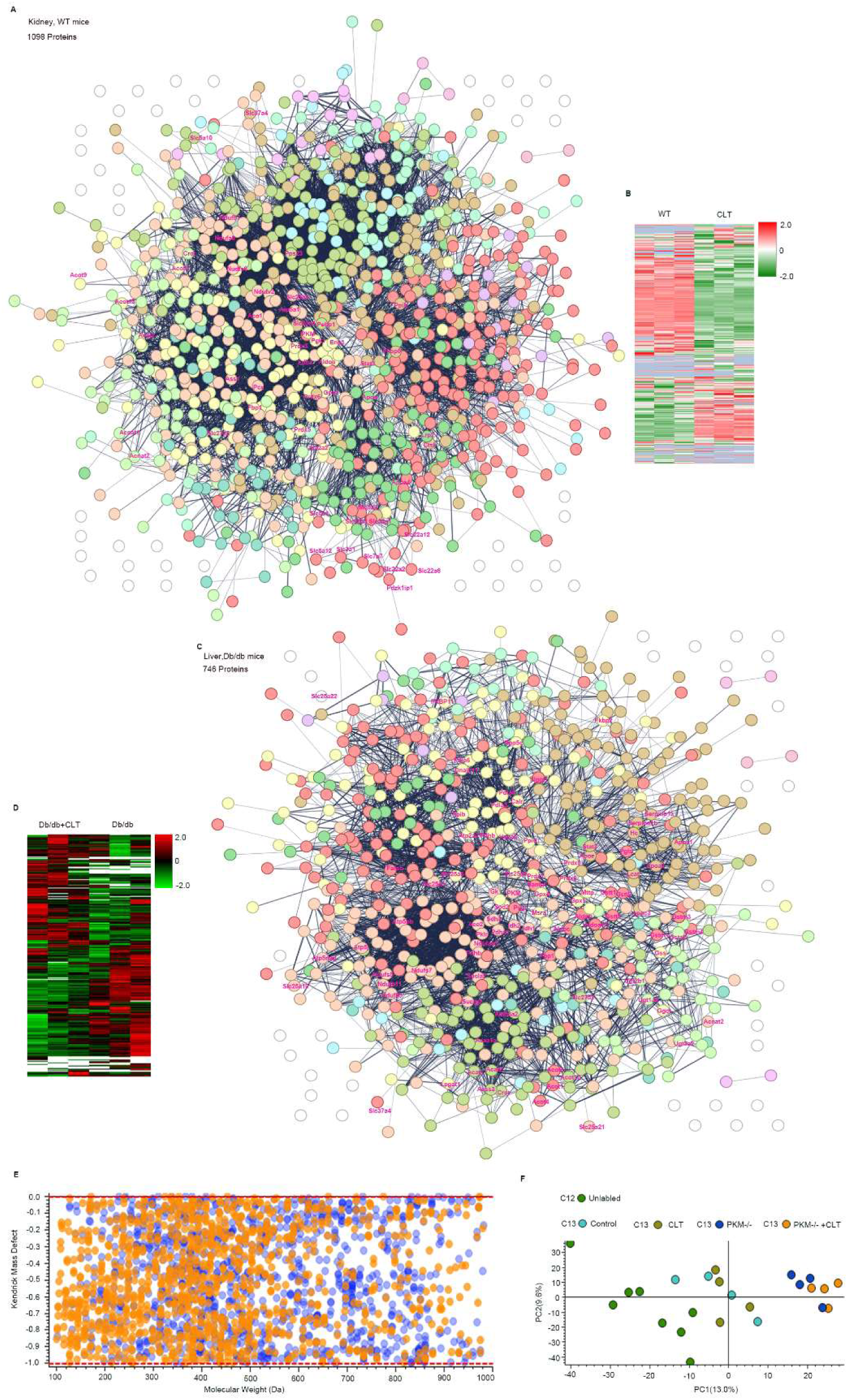
PKM is the centroid of the PPI network of CLT-treated WT mice kidney and CLT-treated mice db/db liver. (A) PPI cluster network of 1023 significantly affected proteins in the kidneys of WT mice after 3 weeks of CLT treatment. n=3, the number of experiments also represents all the following WT mouse kidney proteomics experiment numbers. (B) Proteomics heatmap of 3-week CLT-treated WT mouse kidney. (C) PPI cluster network of 713 significantly affected proteins in 4-week CLT-treated B6-db/db mouse liver. n=3, the number of experiments also represents all the following B6-db/db mouse liver proteomics experiment numbers. (D) Proteomics heatmap of 4-week CLT-treated db/db mouse liver. (E&F) Kendrick mass defect and dotplot of metabolomics sample in unlabeled cells, ^13^C glucose labeled cells, CLT-treated ^13^C glucose labeled cells, PKM-/- ^13^C glucose labeled cells, CLT-treated PKM-/- ^13^C glucose labeled cells. n=4 for ^13^C glucose labeled cell groups, n=8 for unlabeled group, the number of experiments also represents all the following metabolomics experiment numbers.

**Figure 5.**
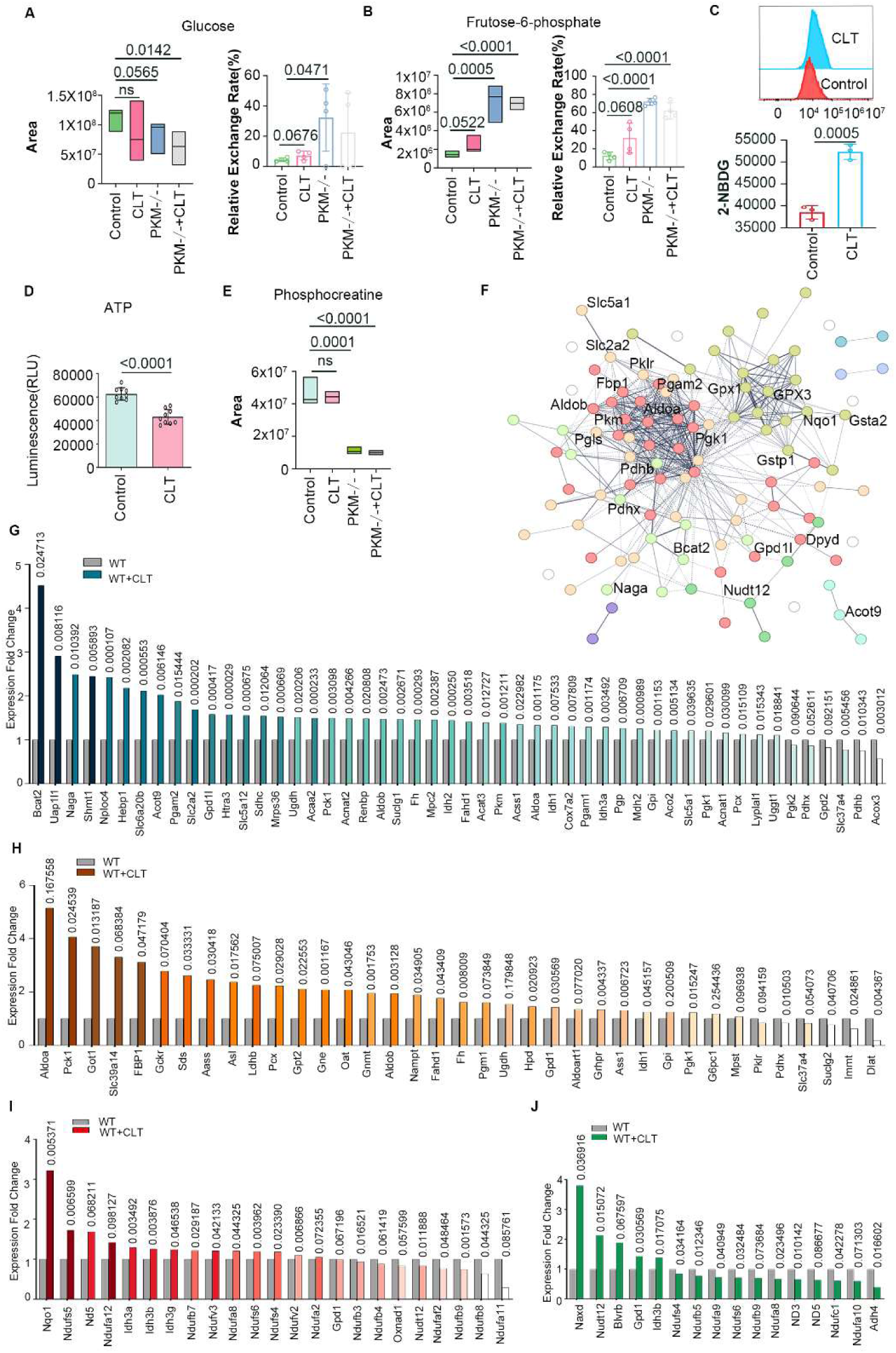
CLT increases the cellular glucose uptake. (A) Glucose levels and ^13^C glucose exchange rates in Control, CLT, PKM -/-, and PKM -/- + CLT A549 cells. Glucose levels decrease in cells while their ^13^C exchange rates increase in response to CLT and PKM knockout treatments. (B) Frutose-6-phosphate levels and ^13^C glucose exchange rates in Control, CLT, PKM -/- and PKM -/- +CLT A549 cells. Both fructose- 6-phosphate and its ^13^C exchange rate increase in response to CLT or PKM knockout treatment. (C) CLT increases the 2-NBDG uptake in HK-2 cells. n=3, p=0.0005. (D) CLT decreases the ATP production in HK-2 cell. n=9, p<0.0001. (E) Phosphocreatine levels decrease in CLT, PKM-/-, PKM-/- +CLT cells. (F) Subcluster of total PPI network cluster in Fig.4A containing PKM. (G) Expression fold change of proteins related to glucose and glycolysis in 3-week CLT-treated WT mouse kidney. (H) Expression fold change of proteins related to glucose and glycolysis in 3-week CLT-treated WT mouse liver, n=3, the number of experiments also represents all the following WT mouse liver proteomics experiment numbers. (I) Expression fold change of proteins related to NADH in 3-week CLT-treated WT mouse kidney. (J) Expression fold change of proteins related to NADH in 3-week CLT-treated WT mouse liver.

To elucidate the alteration of glycolysis, glucose metabolism and energy metabolism signaling pathways, key PPI including PKM, Aldob, Slc5a1, Slc2a2, Fbp1, Pgls, Gpx1, Gpx3 and Nqo1 are shown in Figure 5F. Key protein expression changes upon CLT treatment in WT mice in glycolysis and those glucose related proteins in kidney and liver are also shown in Figures 5G and 5H. Additionally, NADH-related and ATP-related protein expression changes following CLT treatment are presented (Figures 5I and 5J, Figures S4A and S4B). Interestingly, significant protein increases were observed in kidney, including Bcat2, Slc2a2, Slc5a12, Pck1, Aldob, PKM, Aldoa, Pgp, Gpi, Slc5a1, Pgk1 and Pcx. Liver proteins such as Pck1, Fbp1, Pcx, Aldob, Pgm1, Aldoart1, Gpd1, Gpi, Pgk1, G6pc1 are also significantly increased. These results indicate significant changes in the glucose metabolism related to glycolysis or gluconeogenesis. In particular, facilitated glucose transporter member 2 (Slc2a2 / GLUT2) at the basolateral side is positioned to take glucose out of the cell toward the blood (Klip et al., 2024). Therefore, the increased expression of Slc2a2 (Figure 5G), the surge of urine glucose (Figures 3L, 3Q and 3R) and the increasing trend of glucose in cell culture medium (Figure 3S) represents the increase of the cellular glucose outflow. However, Pdhx and Pdhb, which comprise the pyruvate dehydrogenase complex, are both significantly decreased in kidney (Figure 5G). And Phdx is also decreased in liver by CLT (Figure 5H). Since the pyruvate dehydrogenase complex functions following PKM and preparing for TCA cycle, these results imply that the block of PKM might exert possible influence on TCA cycle.

### 4. The inhibition of PKM forms TCA vortex while accelerating the amino acid and lipid degradation as the energy compensation

Cellular metabolism utilizes glucose, amino acids and fatty acids as their source in the TCA cycle (Liu et al., 2020). To examine the TCA cycle metabolomics changes in response to CLT treatment and PKM knockout, the A549 cell extracts were detected by target combined with non-target metabolic LC-MS (Figures 6A-E). Primarily, the citric acid and isocitric acid at the beginning of TCA cycle were both decreased in CLT-treated, PKM-/- and CLT-treated PKM-/- cells (Figures 6A and 6B). Meanwhile, Lys, Leu and tryptophan (Trp) were all decreased (Figures 6F-H) to compensate this shortage (Guertin and Wellen, 2023). However, the downregulation of succinic acid in CLT-treated, PKM-/- and CLT-treated PKM-/- cells is less obvious (Figure 6C). Particularly, His and Pro transform to Glu and supply the above shortage (Jayaraman et al., 2022). In addition, Met, Val, Ile and threonine (Thr) could also supply the TCA shortage (Hara et al., 2021).Therefore, the downregulation of His, Pro, Met, Val, Ile and Thr were detected in CLT treated and PKM -/- or CLT treated PKM -/- cells (Figures 6I-N). In contrast, at the end of TCA cycle, fumaric acid and malate levels increased in CLT-treated and PKM -/- and CLT-treated PKM -/- cells (Figures 6D and 6E). This might due to the further supply of Phe, Tyr, aspartate (Asp) and Asn (Owen et al., 2002), which all decreased in CLT-treated, PKM-/- and CLT-treated PKM-/- cells (Figures 6O-R). The shortage of carbon source caused by PKM inhibition is directly compensated by cathepsin L promoted amino acid catabolism (Figure 6S). Hence, the TCA cycle forms a vortex in response to PKM inhibition.

**Figure 6.**
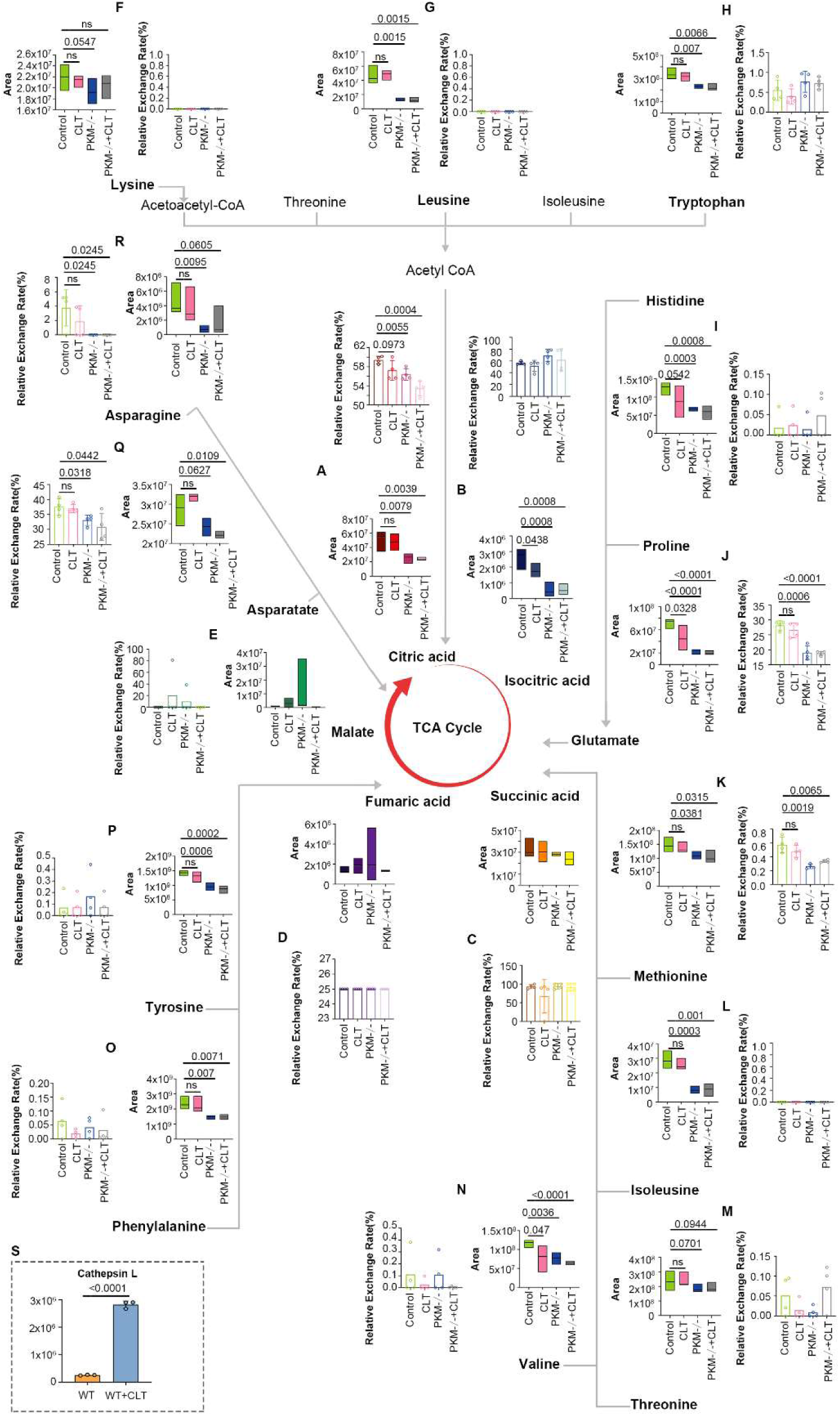
CLT treatment and PKM knockout promote TCA vortex with amino acid degradation as compensation. (A-E) Citric acid, isocitric acid, succinic acid, fumaric acid, malate and their ^13^C glucose exchange rate in the TCA cycle. (F-R) Lys, Leu, Trp, His, Pro, Met, Ile, Thr, Val, Phe, Tyr, Asp, Asn and their ^13^C glucose exchange rates. (S) CLT increased cathepsin L protein expression in WT mice kidney. P<0.0001, n=3.

We next examined the impact of TCA vortex on lipid metabolism. Palmitoylcarnitine, propionylcarnitine, carnitine and acetyl-L-carnitine, 2-hydroxypropyl stearate levels all decreased in CLT, PKM-/- and PKM-/- + CLT group (Figures S4C-I). Lipid acids such as stearate also decreased, though not significant (Figure S4J). On the contrary, glycerophosphocholine and its ^13^C exchange rate were increased by CLT and PKM knockout (Figures S4K and S4L). Therefore, the block of PKM should transfer the glucose carbon flux partially to glycerol metabolism. Followed by the lipid and amino acid degradation, the body weight loss was observed in both 3-week CLT-treated C57BL/6J WT mice and 4-week CLT-treated C57BL/6J B6-db/db mice (Figure S4M). Figure S4N shows the network of lipid metabolism regulating proteins such as Acat2, Acnat1, Acnat2, Crat and Eci3. These proteins together with Acot2, Acot1, Apoe, Lrp1, Lrp2, Apoc3, Fabp4, Fabp5, Acox2, Apoa1, Pisd, and Lcat were either collectively or individually increased in 3-week CLT-treated C57BL/6J WT mice (Figures S4O and S4P). Altogether, the lipid and amino acid degradation compensates for the TCA vortex induced by PKM inhibition.

### 5. Inhibition of PKM shifts the glucose flux and enhances the pentose phosphate pathway (PPP)

To investigate how CLT or PKM knockout affects the glucose flux, the PPP was analyzed as it operates in parallel to upper glycolysis to produce ribose 5-phosphate and nicotinamide adenine dinucleotide phosphate (NADPH) (Teslaa et al., 2023). Remarkably, the levels of PPP closely-related glutathione (GSH) increased in PKM-/- and PKM-/- + CLT, whereas their ^13^C glucose exchange rate was reduced (Figures S5A and S5B). Meanwhile, the levels of important GSH ingredients including glutamate, glutamine, gamma-glutamylcysteine and their ^13^C glucose exchange rate decreased in CLT (not significant), PKM-/- and PKM-/- + CLT (Figures S5C-H). Interestingly, the indirect GSH ingredient cysteinyl-glycine decreased in CLT and PKM-/- (p=0.0771, 0.0768 respectively), while showing bare ^13^C glucose exchange (Figures S5I and S5J). Significantly, direct PPP product 6-deoxy-5-ketofructose 1-phosphate and its ^13^C glucose exchange rate were all increased in CLT, PKM-/- and PKM-/- + CLT (Figures S5K and S5L), indicating enhanced glucose flux into PPP due to PKM inhibition. The network of GSH and its ingredient (Veeravalli et al., 2011) is shown in Figure S5M, where the green indicates reduction and the red means upregulation. The subcluster of Figure 4A related to GSH modulation is shown in Figure S5N. Furthermore, kidney and liver proteomics analyses related to PPP, NADPH and GSH are shown in Figures S5O and S5P. Remarkably, the key PPP enzyme 6-phosphogluconolactonase (Pgls) exhibited a 9.6-fold increase (p=0.036325) upon 3-week CLT treatment (Figure S5O). Taking together, these results confirm that inhibition of PKM by CLT or PKM knockout could shift the glucose flux from glycolysis to PPP, resulting in increased GSH production and enhanced cellular anti-oxidative capability.

### 6. CLT enhances GPX and GSH production and suppresses macrophage inflammatory response

To further understand the potential mechanism of clinical efficacy, we set to examine the anti-inflammation effect of CLT by single cell RNA sequencing (scRNA-seq) in unilateral ureteral obstruction (UUO) kidney with typical aseptic inflammation (Wyczanska and Lange-Sperandio, 2020) and oxidative stress (Dendooven et al., 2011). The scheme of study is shown in Figure 7A. We isolated and sequenced a total of 43,547 cells from whole kidney cell suspensions derived from four healthy male mice (one kidney per mouse). 20,918 cells from CLT group mice kidney and 22,629 cells from UUO group mice kidney were analyzed, respectively. Through data processing and cell type annotation, totally 13 cell types were annotated in our scRNA-seq data (Figure 7B), including tissue-associated cells and immune cells (Park et al., 2018). The tissue-associated cells include podocyte (Podo), fibroblast (Fibro), endothelial and vascular cell (Endo), descending and ascending loop of Henle (LOH), distal convoluted tubule (DCT), proximal tubule (PT), cortical cell (Cor), collecting duct principal cell, collecting duct intercalated cell and collecting duct transitional cell (CD). The immune cells include B cell (B), natural killer/T cell (NK/T), neutrophil (Neutro), monocyte (Mono), and macrophage (Macro). The major marker genes for each cell type are shown in Figure S7A. Remarkably, the immune cell types are the dominant cells in UUO kidney sample, while CLT could significantly eliminate these inflammatory cells and protect the tissue-related cells from being damaged (Figure 7C). Interestingly, the percentage of macrophage is as high as 45%, while CLT treatment could reduce this percentage to 9.81% and partly rescue the tissue-related cells (Figure S7B). We further validated the decreasing number of macrophages in UUO kidney after CLT treatment by immunofluorescent staining (Figure 7D).

**Figure 7.**
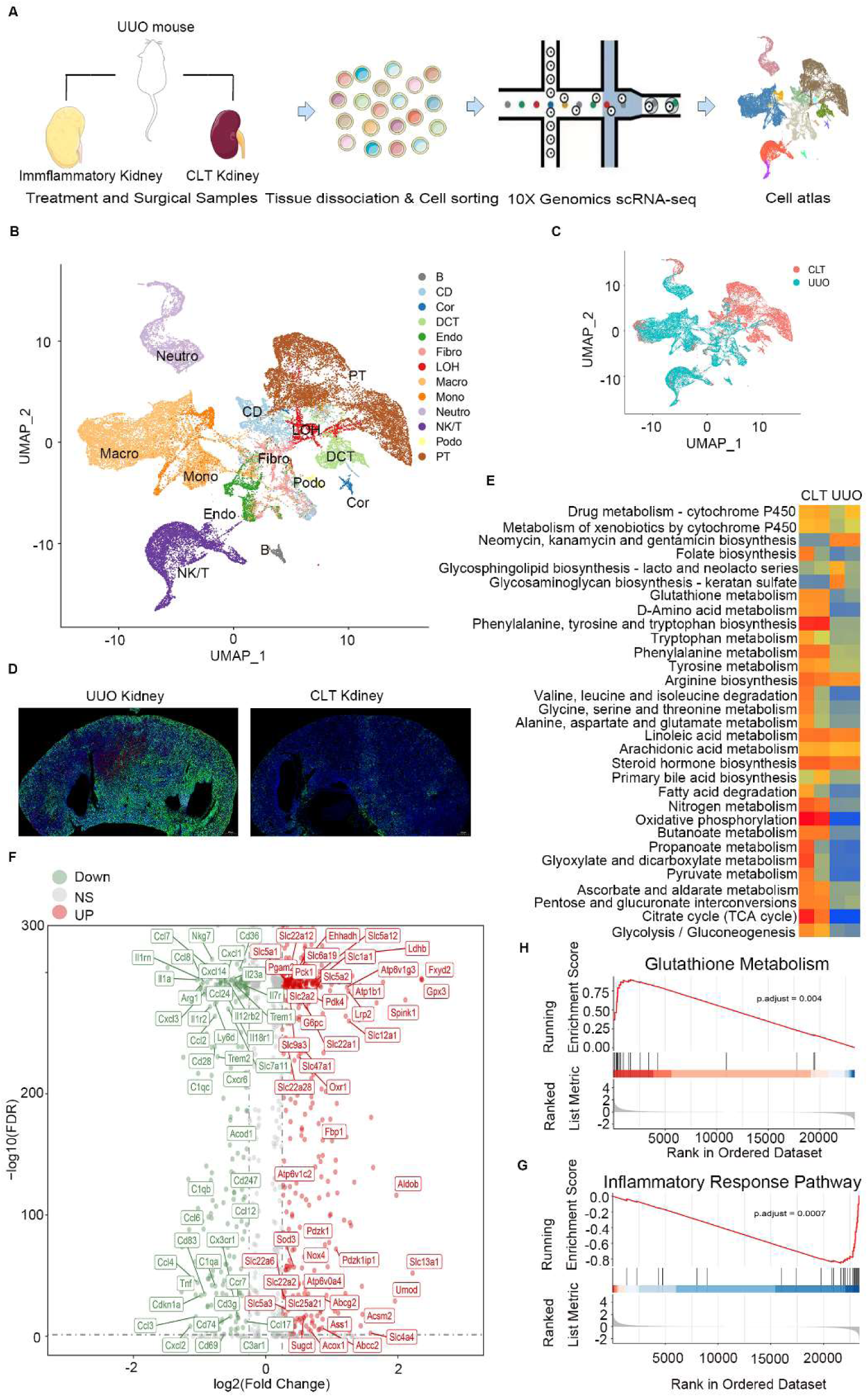
CLT mediates anti-inflammatory effects by inhibiting macrophages. (A) Scheme of scRNA-seq for UUO kidney (n=2) and CLT-treated UUO kidney (n=2). (B) Overall UMAP showing cell annotation from scRNA-seq data. A total of 43,547 cells from four whole kidney cell suspensions were isolated and sequenced. (C) Overall UMAP shown by sample type. 22,629 cells from UUO group mouse kidney; 20,918 cells from CLT group mouse kidney. (D) Immunostaining macrophages by F4/80 (green) in UUO and CLT-treated UUO kidney. The blue indicates DAPI staining. (E) Heatmap showing the comparison of GSVA metabolism-related pathways between CLT-treated UUO kidney and UUO kidney. (F) Volcano map showing 1022 significant DEGs after CLT treatment in UUO kidney. (H) Upregulated glutathione metabolism upon CLT treatment in UUO kidney by GSEA analysis, p=0.004. (G) Downregulated inflammatory response pathway upon CLT treatment in UUO kidney by GSEA analysis, p=0.0007.

To determine the metabolic differences between UUO and CLT treatment groups through scRNA-seq data, we also performed the gene set variation analysis (GSVA) metabolism analysis and found several metabolic processes being significantly up-regulated which includes but aren’t limited to glutathione metabolism, D-amino acid metabolism, oxidative phosphorylation, pentose and glucuronate interconversions, TCA cycle and glycolysis /gluconeogenesis (Figure 7E). Detailed GSVA metabolism analysis result is shown in Figure S6A, which further validates that those amino acids degradation is globally enhanced by CLT (Figure 6F-R). Additionally, the pathways including fatty acid degradation, fatty acid elongation, fatty acid biosynthesis glycolipid metabolism, biosynthesis of unsaturated fatty acids and linoleic acid metabolism were also found to be upregulated by CLT treatment. Generally, these results are highly consistent with the metabolic and proteomics analysis of amino acids and lipid in Figures 5, 6, S4 and S5.

We next investigated the transcriptional differences between the UUO and UUO+CLT groups through differentially expressed gene (DEG) analysis. A total of 1022 DEGs were identified after CLT treatment, among which GPX3 ranked first with a 54-fold upregulation (Figure 7F). Some other important upregulated genes include Slc5a1, Slc5a2, Slc5a12, Slc2a2, Slc1a, Lrp12, Aldob, Pdzk1, Fbp1, G6pc and Sod3, which are consistent with the proteomic results aforementioned. While genes from Cxcl family, Ccl family, Il family, Cxcr family and Cd Family are globally downregulated, indicating that inflammation in UUO kidney are significantly inhibited. To investigate the functional mechanism of CLT treatment, we further performed GSVA analysis and found that the glutathione metabolism related pathway is upregulated, while the inflammatory response pathway is downregulated after CLT treatment (Figures 7H and 7G). Notably, GPX3 was found to be upregulated in all cell types after CLT treatment (Figures S7C and S7D). Moreover, GSVA reactome analysis shows that several genes (i.e. Gpx4-2, Gpx1/2/4, and Gpx3) related to GSH reactions are the major upregulated reactome in CLT treated kidney (Figure S6B), which strongly supports that CLT treatment enhances the anti-oxidative and anti-inflammatory ability by upregulating GSH producing PPP pathway. Further experiments confirmed that NADH and GSH production were upregulated in A549 cells by CLT and PKM knockout, while the ROS level was decreased in these cells. (Figures S7E-I). Taking together, our results reveal CLT targets PKM and exhibit strong anti-oxidative and anti-inflammation capability by shifting the glucose flux from glycolysis to PPP pathway and upregulating the GST system.

## Discussion

### Quantum calculation plus Mass Spectrometer Confirms the CLT-PKM interaction

The quantum computational chemistry and physics calculation (Guo et al., 2024; Santagati et al., 2024) together with the high-resolution, accurate-mass LC-MS system and molecular docking were applied in identifying drug-target interaction. Firstly, DFT calculation perfectly and precisely predicted the interaction patterns of CLT with 20 amino acids (Figures 1, S1 and S2). This could be a versatile approach to explore the possible targets of candidate drugs, especially for those who could form hydrogen bond with amino acids. The vast application of this method combination would hopefully accelerate the discovery of new drug targets. Secondly, the high-resolution, accurate-mass LC-MS identifies PKM as the target of CLT (Figure 2A), consistent with quantum chemistry and physics calculation. Moreover, molecular docking using the Maestro simulation further confirmed the CLT’s binding pocket is the place where ATP is generated from PEP plus ADP (Figures 2C-F). In addition, surface analysis has shown that the binding pocket is constructed by aliphatic amino acids, which allows the binding of liposoluble chemicals such as CLT (Figure S3F). Particularly, string analysis shows that PKM centers the significant protein-protein interaction network (Figures 4A and 4C). This further proves that PKM is the target of CLT. These methodologies, together with multiple omics including transcriptome sequencing, single cell RNA sequencing, metabolomics, and proteomics form an efficient and powerful drug target digging system, rigorously clarifying the CLT-PKM interaction (Figures 5, 6, S4, S5 and S7). This system could be further tested for other small molecule compounds especially for those plant extracts or herbs.

Besides, the CLT binding pocket is an physiological pocket for ATP (PDB: 4FXF)(Morgan et al., 2013) and pyruvate (PDB:4YJ5) (Chen et al., 2019). Both CLT and ATP could inhibit PKM enzyme activity (Figure 2I) (Callens et al., 1991). Basically, CLT acts as a stiff “ATP” with a similar size which imitates the binding to PKM of ATP and inhibits the enzyme activity. CLT shows longer and stronger inhibition to PKM probably because CLT has a stable structure and does not hydrolyze and release phosphate like ATP (Fontecilla-Camps, 2022). Thus, it is reasonable to speculate that the CLT-PKM interaction could be a perfect reflection for ATP-PKM interaction in regulating glucose metabolism and energy expenditure. When this same pocket is occupied by CLT, the natural glucose metabolism and energy corresponding to ATP’s PKM inhibition is activated. Therefore, CLT’s unimaginable pharmacological and physiological effect on glucose, obesity and inflammation should reasonably root from some existing while undiscovered energy and glucose regulatory mechanism for ATP.

### PKM inhibition induced glucose carbon flux U turn and TCA vortex

The inhibition of PKM by CLT results in two major phenomena that appear contradictory: the inhibition of glycolysis and the decrease in blood glucose. Unexpectedly, the uptake of the glucose (Figures 5A and 5C) is consistent with the downregulation of blood glucose (Figures 3E-G), but contradictory to the glycolysis inhibition (Figures 3A-D). We set to uncover the underlying mechanisms in the downstream and other closely related metabolic pathway, including TCA cycle, PPP, amino acid and lipid metabolism (Martínez-Reyes and Chandel, 2020).

The glucose U turn is corresponding to the PKM enzyme activity as well as the TCA vortex. Normally, the inhibition of PKM (Figure 2I) could lead to the decrease of glycolytic flux (Figures 3A-D) and then the less uptake of glucose. In contrast, both the glucose uptake and its ^13^C exchanging rates increase in response to PKM inhibition (Figures 5A and 5C). In addition, the glycolytic inhibition starves the cell, resulting in low ATP and phosphocreatine state (Figures 5D and 5E). And this continuous shortage of glucose and energy conduce to TCA cycle imbalanced cycling, shaping TCA cycle as a vortex (Figures 6A-E). Typically, the TCA vortex induced by glucose-shortage promotes the amino acid degradation (Figures 6F-R) and lipid degradation (Figures S4C-J) as forms of energy compensation. Besides, the important product of glucose for glycolysis frutose-6-phosphate accumulates significantly (Figure 5B) but could not produce energy in normal glycolytic -TCA flux. Instead, the glucose carbon flux makes a U turn in front of PKM (Graphic abstract), flowing to other fluxes such as PPP (Figures S5K and S5L).Meanwhile, other glucose-sourced metabolites such as glycerol production is increased as well (Figures S4K and S4L), showing a contrary trend comparing to other lipid metabolites (Figures S4C-J). Hence, the TCA vortex and ATP-shortage must be the major driving force to uptake more glucose, though the clear signaling needs to be further mapped. Consequently, our data solidly validate the anti-obesity effect of CLT (Figure S4M) (Liu et al., 2015; Ma et al., 2015).

However, what’s the destination of the increased glucose inflow? One important split-flow of glucose carbon U turn flux flows to the PPP. The levels of 6-deoxy-5-ketofructose 1-phosphate and its ^13^C glucose exchange rate increased following CLT and PKM knockout (Figures S5K and S5L), producing abundant GSH (Figures S5A and S5B). Similarly, the inhibition of PKM by ATP stimulates the PPP pathway to generate more GSH, combatting the oxidative damage when oxidative phosphorylation is active (Stincone et al., 2015; Teslaa et al., 2023). In fact, CLT exhibits extraordinary anti-oxidative and anti-inflammation capacity (Hu et al., 2017; You et al., 2021). Thus, the inflammatory cell number and signals declined following CLT treatment, while all the glycolytic changes could be clearly seen by single cell analysis (Figures 6B-G, Figure 7B). Especially, GPX3 gene ranks first among all increasing responding genes (Figure 6F). Remarkably, CLT-PKM interaction induced GPX enhancement plays a crucial role in the anti-inflammation response (Figure 6H, Figure 7D and Figure S7B). Moreover, some other natural compounds could increase glutathione levels while the action mechanism is unknown (Di Giacomo et al., 2023). Interestingly, ATP may exhibit anti-inflammatory properties with prolonged exposure (Faas et al., 2017; Orimoto et al., 2024). Hence, the anti-inflammatory capabilities of CLT and ATP likely share a common mechanism, inhibiting PKM and enhancing PPP. Subsequently, additional experiments are needed to investigate the interactions of natural compounds with PKM.

### ATP resistance other than insulin resistance in diabetes pathogenesis

CLT-PKM interaction vividly reproduces the ATP-PKM interaction during diabetes mellitus. Importantly, CLT binds to exactly the same site on PKM as ATP (Figures 2C-H). In addition, this binding leads to stronger inhibition of PKM enzyme activity than that of ATP inhibition (Figures 2F and 2I). Therefore, the CLT-PKM interaction and the corresponding effect could reflect the ATP-PKM interaction in the biochemical and physiological process of diabetes.

Many remarkable gaps in diabetes and glucose metabolism still remain unknown in current knowledge (Klip et al., 2024). However, very recent attention has noticed that glycolysis as well as PKM has high control strength over insulin secretion (Merrins et al., 2024). Meanwhile, the accumulation of fat, protein, and sugar results in oxidative phosphorylation-driven rise in ATP (Merrins et al., 2024). The increased ATP could then inhibit PKM (Callens et al., 1991), and enhanced gluconeogenesis is also a significant contributor to hyperglycemia in diabetes mellitus (Prié, 2014 ; Barroso et al., 2024). Remarkably, CLT reproduced these processes including increased urine glucose excretion (Figure 3R) and enhanced gluconeogenesis (Figure 6E). Nevertheless, what is the physiological and pathological principle between hyperglycemia and weight-loss for diabetes mellitus? Why high blood glucose could lead to the loss of body weight? Cellular ATP-PKM interaction imitated by CLT-PKM might be a perfect interpretation for these gaps.

Accordingly, glucose efflux is another important split-flow of the block of glucose flux by CLT-PKM interaction. The blocked glycolysis metabolism pathway by CLT or PKM -/- treatment leads to the piled up glucose and glycolysis pathway goes more to the gluconeogenesis side (Figures 6E and S7A), thereby forming a carbon flux U turn just in front of PKM (Graphic abstract). Hence, we saw the increasing of gluconeogenesis and the increased excretion of glucose (Figure 3S). Notably, a significant urine glucose increase was detected on the first day of CLT treatment (Figure 3R), consistent with the bladder observation 18F-FDG signal of PET/CT (Figure 3J). Moreover, enhanced bladder signals of 18F-FDG in PET/CT scans (Figures 3I-K) indicate increased urine glucose. In addition, a 1.68 fold increase in Slc2a2/GLUT2 protein expression (Figure 5G) and 249 fold increase of glucose-6-phosphatase gene expression (Bulk RNA sequencing data) were detected. Particularly, Slc2a2/GLUT2 at the basolateral side is positioned to take glucose out of the cell toward the blood (Klip et al., 2024; Leturque et al., 2009). Additionally, glucose-6-phosphatase could convert glucose-6-phosphate to glucose (Klip et al., 2024). Taking together, all these data support that inhibition of PKM by CLT could accelerate the glucose outflow by upregulating gluconeogenesis as well as Slc2a2. Consistently, in muscles where lacking glucose-6-phosphatase (Jensen et al., 2011), the decreased muscle 18F-FDG intake in 21-day CLT treated mice and PKM knockout mice was observed (Figures 3J, 3K and 3O). These finding strongly indicate that the PKM inhibition effect on glucose U turn depends on full gluconeogenesis chain including glucose-6-phosphatase. Therefore, increased 18F-FDG uptake could occur in tissues expressing glucose-6-phosphatase (Figure 3O). Consequently, the increased cellular uptake of glucose (Figures 5A and 5C) and the enhanced glucose efflux collectively contribute to the reduction in blood glucose levels (Figures 3E-G).

In summary, the CLT-PKM interaction could be an ideal model for ATP-PKM interaction in diabetes mellitus. Continuous inhibition of PKM by ATP during the development of diabetes leads to glycolysis inhibition. This, in turn, altered the glucose carbon flow and caused hyperglycemia, resulting in hyperglycemia and glycosuria (Prié, 2014). Hence, ATP-PKM interaction induced TCA energy vortex causes weight loss, which has been reproduced by CLT-PKM interaction, as an important symptom of diabetes. The inhibition of PKM by ATP is a chronic process that accumulates and persists over time. This weak and short inhibition (Figure 2I) might be a protective reaction of the body, leading to the increase of the blood glucose (Figures 3F and 3J). However, CLT is a long and strong inhibition (Figure 2I), which amplifies and accelerates the inhibitory effect of ATP, immediately reducing the blood glucose levels (Figure 3J) by increasing the cellular glucose uptake (Figures 5A and 5C) and enhancing the urine glucose excretion (Figures 3L and 3R). Thus, CLT functions to enhance and amplify ATP’s inhibitory effect on PKM. Our data indicate “ATP resistance” at the cellular level for the diabetes in addition to the “insulin resistance”. More effort should be spent to directly investigate ATP’s inhibition of PKM during the pathogenesis of diabetes. Meanwhile, the inhibition of PKM by CLT to treat diabetes provides a new target for the science community, offering a more powerful tool to control the increasing obesity and diabetes.

### Limitations of the study

It should also be noticed that in this DFT simulation, we do not consider all the possible interaction patterns between CLT and amino acids. Therefore, our results might be modified and improved by further complete simulation. We do not specifically study the detail how glucose-6-phosphatase cooperates with Slc2a2 in transporting glucose into the bloodstream. Further rigorous experiments are needed in the future to explain the precise molecular and biochemical mechanisms by which glucose exhibits hyperglycemia and hypoglycemia in ATP low-intensity inhibition and CLT high-intensity inhibition, respectively. Since ATP is easily hydrolyzed, a critical challenge is to develop appropriate model and technique to directly map the ATP-PKM interaction in the glucose metabolism and diabetes pathogenesis.

## Acknowledgments

This work is supported by the Natural Science Foundation of Chongqing (CSTB2023NSCQ-MSX0620, cstc2021jcyj-msxmX0946, cstc2019jcyj-msxmX0408), the Technology Foresight and Institutional Innovation Project of Chongqing (CSTB2023TFII-DIX0051), the National Natural Science Foundation of China (81901111) & the Open Project of Key Laboratory of Electromagnetic Radiation Protection, Ministry of Education, China (2022DCKF). We thank Dr. Renxiao Wang & Dr. Yan Li for assistance in computer resources provided by the School of Pharmacy, Fudan University. Dr. Xiaoyang Wang, Dr. Feng Pan & Dr. Wenjun Shan provide technique assistance in proteomics, metabolomics and animal experiments. Dr. Zeping Hu & Dr. Bohong Wang from Tsinghua University give useful suggetion in metabolomics experiment design and result analysis. Dr. Mindian Li at Southwest Hospital, Third Military Medical University kindly provides the Seahorse XF mitochondrial stress and glycolysis stress test kits. Dr. Zongshi Lu at Daping Hospital, Third Military Medical University kindly helps us collect and analyze the 18F-FDG PET/CT imaging data. Dr. Xiaojin Pan & Dr. Qifan Wang at Tsinghua University generously provide the pyruvate kinase PKM recombinant protein.

## Author contributions

Conceptualization, Q.C. and X.Z.; Methodology, Q.C., X.Z., X.D.J., Y.H.Y., J.T.W., and M.Z.; Validation, Q.C. and X.D.J.; Formal analysis, Q.C., X.D.J., X.Z., X.B.W., and Y.H.Y.; Investigation, Q.C., X.D.J., X.Z., Y.H.Y., and J.F.Z.; Data curation, Q.C., X.Z., X.D.J., Y.H.Y., and X.B.W.; Resources, Q.C., M.Z., and X.B.W.; Writing – original draft, Q.C., X.Z., and Y.H.Y.; Writing – review & editing: Q.C., Y.H.Y, X.B.W., and M.Z.; Visualization, Q.C and X.Z.; Supervision & Funding acquisition, Q.C and X.Z.

## Methods

## KEY RESOURCES TABLE

| REAGENT or RESOURCE | SOURCE | IDENTIFIER |
| --- | --- | --- |
| **Antibodies** |  |  |
| Rabbit monoclonal [EPR10138(B)] to PKM | Abcam | Cat# ab150377, RRID:AB_2941747 |
| Mouse monoclonal [mAbcam 8226] to beta Actin | Abcam | Cat# ab8226, RRID:AB_306371 |
| **Chemicals, peptides, and recombinant proteins** |  |  |
| D-glucose-<sup>13</sup>C<sub>6</sub> | MedChemexpress | HY-B0389A<br>CAS:110187-42-3 |
| D-glucose | TargetMol | T0887 |
| Fumaric acid | TargetMol | T3763<br>CAS:110-17-8 |
| Malate | TargetMol | T2S0850<br>CAS:6915-15-7 |
| DL-isocitric acid trisodium salt | TargetMol | T5087 |
| D-glucose 6-phosphate sodium | TargetMol | T4726<br>CAS: 54010-71-8 |
| Fructose-6-phosphate | TargetMol | T4941<br>CAS: 26177-86-6 |
| Succinic acid | TargetMol | T2719 CAS:110-15-6 |
| Citric acid | TargetMol | T5S0636 CAS:77-92-9 |
| Itaconic acid | TargetMol | T4837 CAS:97-65-4 |
| Seahorse XF DMEM | Agilent Technology | 103575-100 |
| 18F-FDG | Zhiming Zhu(Daping Hospital, Army Medical University) | N/A |
| Streptozotocin | Aladin | S110910 |
| Celastrol | Sichuan Vicky Biotechnology | WKQ-0000332<br>CAS:34157-83-0 |
| 2-NBDG | CSNPHARM | A422362<br>CAS:186689-07-6 |
| PKM peptide | Xiaojing Pan(Tsinghua University) | N/A |
| Adenosine 5'-triphosphate disodium salt hydrate | Sigma-aldrich | A26209<br>CAS:34369-07-8 |
| **Critical commercial assays** |  |  |
| Pyruvate kinase assay kit | Sigma-aldrich | MAK072 |
| Mouse glucose assay kit | Crystal Chem | Cat# 81692 |
| Enhanced ATP assay kit | Beyotime Biotechnology | S0027 |
| Reactive oxygen species assay kit | Beyotime Biotechnology | S0033S |
| NADP+/NADPH assay kit with WST-8 | Beyotime Biotechnology | S0179 |
| **Deposited data** |  |  |
| Sequence data would be deposited online after acceptance | This paper | XXXXX |
| Metabolomics data would be deposited online after acceptance | This paper | XXXXX |
| Proteomics data would be deposited online after acceptance | This paper | XXXXX |
| **Experimental models: Cell lines** |  |  |
| Human A549 cell line | Cyagen | Cat# CM01B005<br>https://www.cyagen.com/cn/zh-cn/business/tumor-cell-list/WT/A549.html |
| Human PKM knockout A549 cell line | Cyagen | SY-KO-00010<br>https://www.cyagen.com/cn/zh-cn/cell-bank-cn/SY-KO-00010 |
| Human HK-2 cell line | National Collection of Authenticated Cell Cultures | SCSP-511<br>https://www.cellbank.org.cn/search-detail.php?id=504 |
| Human PKM knockout HK-2 cell line | Cyagen | SY-KO-00465<br>https://www.cyagen.com/cn/zh-cn/cell-bank-cn/SY-KO-00465 |
| **Experimental models: Organisms/strains** |  |  |
| Mouse: C57BL/6J B6-db/db | Cyagen | C001291<br>https://www.cyagen.com/cn/zh-cn/business/drug-filter-evaluate-mouse/metabolism-medicinal.html |
| Mouse: C57BL/6J-Pkm<sup>em1C</sup>/Cya | Cyagen | S-KO-03667<br>NCBI:18746<br>https://www.cyagen.com/cn/zh-cn/sperm-bank-cn/S-KO-03667 |
| Mouse: C57BL/6J | Animal Center of the Third Military Medical University | N/A |
| Mouse: BALB/c | Animal Center of the Third Military Medical University | N/A |
| **Software and algorithms** |  |  |
| Gaussian09 | Frish et al. | https://gaussian.com |
| Vienna Ab-initio Simulation Package(VASP) | Georg Kresse | 6.0 Version<br>https://www.vasp.at/ |
| Gauss View | Frish et al. | 6.0.16 Version<br>https://gaussian.com/gaussview6 |
| Graphpad Prism | Dotmatics et al. | 8.0.2 Version<br>https://www.graphpad.com/scientific-software/prism/ |
| FlowJo | BD Biosciences | 10.8.1 Version<br>https://www.flowjo.com/solutions/flowjo |
| Image Lab | BIO-RAD | 5.2 Version<br>https://www.bio-rad.com/zh-cn/category/image-lab-software-resources?ID=PJWA0VTU86LJ |
| Maestro | Schrödinger | 12.9 Version<br>https://www.schrodinger.com/platform/products/maestro |
| Compound Discoverer | Thermo Fisher Scientific | 3.3.1.111Version<br>https://www.thermofisher.cn/cn/zh/home/industrial/mass-spectrometry/liquid-chromatography-mass-spectrometry-lc-ms/lc-ms-software/multi-omics-data-analysis/compound-discoverer-software.html |
| Thermo Proteome Discoverer | Thermo Fisher Scientific | 2.5.0.400 Version<br>https://www.thermofisher.cn/cn/zh/home/digital-solutions/ardia-platform/proteomics.html |
| MzVault | Thermo Fisher Scientific | 2.1 Version<br>https://www.thermofisher.cn/cn/zh/home/industrial/mass-spectrometry/liquid-chromatography-mass-spectrometry-lc-ms/lc-ms-software/mass-spectral-libraries.html |
| Cellranger Count | 10x Genomics | https://www.10xgenomics.com/support |
| Seurat | Rahul Satija | 4.3.0.1 Version<br>https://satijalab.org/seurat/ |
| Cellscope | CapitalBio Technology | http://101.39.241.142:7683/#/ |
| Avatar | PINGSHENG HEALTHCARE | https://www.pingsheng.com/product/default_1.htm#item |
| Cruiser | PINGSHENG HEALTHCARE | 1.6.8.0 Version<br>https://www.pingsheng.com/product/default_1.htm#item |
| STRING | Damian Szklarczyk etc. | https://cn.string-db.org/cgi/input?sessionId=bHJaskniGY1G&input_page_show_search=on |

### RESOURCE AVAILABILITY

#### Lead contact

Requests for reagents and resources should be directed to and will be fulfilled by the lead contact, Dr. Qian Chen.

#### Materials availability

Reagents generated in this study will be made available by reasonable request to the lead contact.

#### Data and code availability

The accession numbers for data and code would be deposited and listed in the key resources table after acceptance. Other relevant data are available from the corresponding authors upon reasonable request.

### COMPUTATIONAL DETAILS

#### VASP Calculations

First-principles calculations were performed as previously described (Chen et al., 2020; Zheng et al., 2023) within non-spin-polarized density functional theory (DFT) as implemented in the Vienna ab initio simulation package (VASP) (Kresse et al., 1996). The structural optimization and the electronic property were calculated within the generalized gradient approximation (GGA) proposed by Perdew, Burke, Ernzerhof (PBE). Projector augmented wave (PAW) potentials were employed to describe the ion-electron interaction (Blöchl, 1994). We adopted the optPBE-vdW functional to duly treat the effect of the van der Waals interaction between CLT and amino acids. The molecular structures were all obtained from PubChem. The lattice constant of a unit cell was set to 10×10×10 Å, which has a single amino acid in orthorhombic primitive unit cell. In order to avoid the interaction between molecules in neighboring cells, the combination of CLT and amino acids were put in a cell with lattice constant 20×20×20 Å. The initial distance between CLT and amino acid was set to 3 Å. The energy cutoff was set to 400 eV, and the Monkhorst−Pack k-grid mesh was 5×5×1 in surface calculations. Before energy and electronic structure calculation, all atom were fully optimized until the force of each atom was less than 0.03 eV/Å.

The adsorption energy (*E_a_*) between different amino acids with CLT was computed used the following equation:

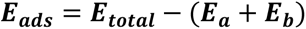

where *E_total_* represents the total energy of the whole combination system. *E_a_* is the energy of CLT and *E_b_* is the energy of isolated amino acids, respectively. By this definition, larger negative value of *E_ads_* denotes stronger interaction between molecules.

#### Gaussian Calculations

The geometry of the molecules considered in this study was preliminarily optimized with DFT as implemented in the VASP with the GGA proposed by PBE. Then the optimized model was calculated by Gaussian 09 software package in the ground state(Cruz et al., 2023). Theoretical calculations of the CLT and amino acids were performed at hybrid B3LYP method using 6-31G basis set. Using Gauss View 6. 0 software (Dixit and Yadav, 2015), the molecular electrostatic potential (MEP) surface of CLT and amino acids were visualized at the ground state energy level without applying any constraint on the potential energy surface. With MEP analysis, the potential interaction sites can be located by different color codes. The red region in a MEP surface map indicates a site tend to gain electrons, while the blue region indicates a site apt to lose electrons. On these structures, the MEP was evaluated and analyzed, and we predicted intermolecular binding sites from the resulting information.

#### Molecular docking

Schrödinger Maestro (v12.9, New York, United States) was used for molecular docking, which consists of various modules including LigPrep (ligand preparation), SiteMap (finding the ligand binding site in the target), GlideGrid (fixing the ligand binding site), and GlideDock (for assessing binding affinities). Ligand CLT was adjusted to a theoretical pH of 7.0, and the ligprep panel within the Maestro software was used. Desalting and tautomer generation methods were applied to optimize calculations while limiting the number of stereoisomer calculations to approximately thirty per ligand and using the output format specified by Maestro software. PKM crystal structures of different packing orders (PDB entries 4FXF, 4YJ5 and 3ME3) were obtained from the RCSB database (http://www.rcsb.org/pdb) and imported into Maestro workspaces. Protein structure (3ME3) was processed using the Protein Production Wizard in the Schrodinger suite. During protein preprocessing, bond orders were assigned, water molecules were removed, and protonation states were adjusted to pH 7.0. Water molecules with fewer than three hydrogen bonds to non-water entities were eliminated. To delineate the operational binding regions for ligand docking tasks, receptor grid files were created using the receptor grid formation panel. The docking score and docking orientation for ligand-protein were documented after molecular docking by glidedock module. Hydrogen bond is shown in the ligand interaction diagram. Binding site surface was created for ligand and surface as solid style with 30% surface transparency. The interaction map is shown in mesh style, with red-while-blue color ramp for the protein while rainbow color ramp for the ligand. The electrostatic potential was set in the range from -0.3 to 0.3.

### EXPERIMENTAL MODEL AND SUBJECT DETAILS

#### Mice

C57BL/6J B6-db/db and C57BL/6J-Pkmem1C/Cya were purchased from Cyagen Biosciences (Suzhou, China). The C57 BL/6J WT mice and BALB/c WT mice were purchased from the Animal Center of the Third Military Medical University (Chongqing, China). All the mice were used for treatment at 6-8 weeks of age, and both the male and female gender mice were included without randomization or“blinding”. All mice were housed in specific-pathogen free (SPF) condition. All mouse experiments and breeding conditions were performed in accordance with the guidelines of the Institutional Animal Care and Use Committees of the Third Military Medical University.

#### WT mouse and design

The healthy C57 BL/6J WT mice were divided into two groups: the control group and the CLT (1mg/kg) injected group, administered intraperitoneally every day. Blood glucose levels and body weights were measured 6 hours after injection. After 3-week treatment, all animals were sacrificed. Their kidneys and livers were collected for proteomics analysis.

#### Type 1 diabetes mouse and design

The type1 diabetes mouse model was carried out as mentioned earlier (Rustagi et al., 2022). STZ was purchased from Aladin (Shanghai, China). STZ was rapidly dissolved in 0.05 mol/L citrate buffer solution with a pH value of 4.5. It was prepared and used immediately on ice, and injection was completed within 30 minutes. C57 BL/6J WT mice were adaptively raised for 7 days, fasting blood glucose levels and body weights data were recorded before modeling (fasting but provided drinking water), followed by intraperitoneal injection of STZ 180mg/kg. 72 hours after injection of STZ, those with blood glucose values > 11.1 mmol/L are considered successful in modeling. Their blood glucose values and body weights data are recorded. High CLT (1mg/kg) treatment and low CLT (0.5mg/kg) treatment were adopted in experiment, while the STZ group was treated with 0.9% NaCl. Intraperitoneal injection was performed every day. Blood glucose levels and body weights were measured 6 hours after injection.

#### B6-db/db mouse and design

C57BL/6J B6-db/db mice were used to test the hypoglycemic effect and anti-obesity of CLT. The B6-db/db mice were divided into two groups: the Control group was injected with 0.9% NaCl and the CLT group was injected with sterile CLT (1mg/kg), administered intraperitoneally every day. Blood glucose levels and body weights were measured 6 hours after injection. After 4-week treatment, all animals were sacrificed. Their tissue was collected for proteomics analysis.

#### PKM knockout mouse and design

C57BL/6J WT mice, CLT injected C57BL/6J WT mice (1mg/kg) and PKM knockout mice (C57BL/6J-Pkmem1C/Cya) were used in the experiment of 18F-FDG PET/CT experiment. Their blood glucose levels were also recorded for 2 weeks.

#### UUO models and design

UUO model was set up to study inflammation and oxidative stress (Dendooven et al., 2011; Wyczanska and Lange-Sperandio, 2020). The UUO model of BALB/c mice (6 ∼ 8weeks) was performed as previously described (Zhang et al., 2021). The mice were divided into two groups: the UUO group was treated with 0.9% NaCl, and UUO + CLT group received CLT (1 mg/kg), administered intraperitoneally every day. The body weights were recorded every two days. All animals were sacrificed on day 14 and their kidneys were harvested for sequencing analysis.

#### Cell lines and design

Human A549 cell line & PKM knockout A549 cell line, Human HK-2 cell line & PKM knockout HK-2 cell line were cultured for different experiments. Briefly, cells were cultured in DMEM/F12 and growth medium containing 10% fetal bovine serum (FBS) in a moist 5% CO_2_ environment, with a constant temperature of 37 ℃. And the cells grew to 85% density before treatment. Then, the cells were rinsed with PBS (1×) and all the liquid was dried before harvest. The cell experiment design is described in separate methodology part.

### EXPERIMENTAL METHOD DETAILS

#### 18F-FDG PET/CT imaging

Glucose uptake in mice was assessed on Super Nova PET/CT scanner (Pingseng Healthcare, Kunshan, China) as previously reported (He et al., 2022; He et al., 2019). On the day of experiment, C57BL/6J WT mice, CLT treated C57BL/6J WT mice (1mg/kg) and PKM knockout mice (C57BL/6J-Pkmem1C/Cya) were anesthetized with isoflurane and injected intravenously with around 8MBq 18F-FDG after fasting for 12 hours. Following a 60 minute absorption period, mice were imaged on the Micro-PET-CT scanner. The CT images were acquired using a 70 kV X-ray tube operating at 600 μA. 10 minute PET images and dynamic 3-dimensional scans were acquired and CT was utilized to obtain anatomical reference images. The PET image was then reconstructed using ordered subsets expectation maximization followed by maximum a posteriori via Avatar software (Pingseng Healthcare, Kunshan, China). Uptake rates of 18F-FDG were analyzed as volumes of interest (VOIs) were drawn over different organs or regions. The rate of glucose utilization was measured using standard uptake value (SUVs) (SUV = [18F-FDG activity in each VOI (Bq/mL)] / [injected dose (Bq)]/ [body weight (g)]).

#### Mouse urine glucose test

Mouse urine glucose was tested using mouse glucose assay kit (Crystal Chem Illinois, USA). Briefly, 2 µL of mouse urine was added to the 300 µL of enzyme solution. And the mixed solution was incubated at 37 °C for 5 minutes. Then, the solution optical density (OD) at 505 nm was measured by SpectraMax i3x (Molecular Devices, Wals, Austria).

#### Immunofluorescence staining

Kidney tissue harvested from animals on day 14 was fixed in 4% paraformaldehyde, embedded in paraffin and cut into 4 µm thick per section. The primary antibodies used were as followed: rabbit anti-OLR1 (1:200, Affinity Biosciences, China), mouse anti-F4/80 (1:200, Abcam, CA, USA). And FITC labeled donkey anti-mouse antibody (1:200, Servicebio, Wuhan, China), Alexa-Fluor 594 labeled goat anti-rabbit antibody (1:400, Jackson, Pennsylvania, USA) were as secondary antibodies. DAPI was used to stain the neucler. Sections and cells were mounted and visualized using Pannoramic MIDI Digital Slide Scanner (3DHISTECH, Budapest, Hungary).

#### Glucose uptake assay

HK-2 cells with 5 × 10^5^ cells/well were laid flat on a 6-well cell plate and cultured in a 5% CO_2_ incubator at 37 °C. After the cell density reached 85%, the drug was administered. The treatment groups were as follows: Blank, Control, CLT. Before administration, the culture medium was replaced with serum-free medium in each group. CLT group was added with CLT (500nM) while control group was added DMSO solution without CLT for 24h. Subsequently, except for the blank group, other two groups were added with 2-NBDG (50 μM) and incubated in the dark for 1 hour. Subsequently, the cells in each well were digested and collected using trypsin, followed by staining with DAPI (500ng/mL) for 3 minutes. After staining, flow cytometry detection was performed on CytoFlex LX Flow Cytometer (Beckman Coulter, California, USA) within 30 minutes. For each test, flow cytometry was used to collect data from 20000 single cells. Data was analyzed by FlowJo (BD Biosciences, New Jersey, USA).

#### Seahorse XF mitochondrial stress and glycolysis stress test

The intracellular mitochondrial stress and glycolysis pressure were detected using the XF96 Extracellular flux Analyzer (Agilent Technologies, Santa Clara, USA) according to the reagent manufacturer’s instructions. Prior to the assay, XF sensor cartridges were hydrated on the day before experiment. To each well of an XF utility plate, 200 uL of sterile water was added; subsequently, the XF sensor cartridges were placed at top of the utility plate, and kept at 37 °C incubator without CO_2_ overnight. Four A549 cell groups were set up: Control, CLT, PKM-/- and PKM-/- +CLT.

For mitochondrial stress test, cells were seeded at 1.5 × 10^4^ cells/well in normal growth media in a Seahorse XF-96 cell culture microplate. The plate was then incubated overnight before drug administration treatment. Basal oxygen consumption rate and spare respiratory capacity were investigated using a Seahorse XF-96 cell mito stress assay according to the manufacturer’s instructions. Following CLT treatment, the drugs and concentrations added in each dosing well were as follows: oligomycin (1.5 μM) in A wells, carbonyl cyanide-4-(trifluoromethoxy)phenylhydrazone (FCCP) (1 μM) in B wells, and rotenone/antimycin A (0.5 μM) in C wells. The different respiration parameters were obtained by subtracting the average respiration rates before and after the addition of the special drugs. The Agilent mitochondrial pressure assay report generator (Agilent Technologies, Santa Clara, USA) was used to analyze the pressure changes of cell mitochondrial in each experimental group.

For glycolysis stress test, cells were seeded at 1.5 × 10^4^ cells/well in normal growth media in a Seahorse XF-96 cell culture microplate. The plate was then incubated overnight before drug administration treatment. The extracellular acidification rate (ECAR) was measured after CLT treatment. Then the drugs and their concentrations added in each dosing well were as follows: glucose (10 mM) in A wells, oligomycin (1 μM) in B wells, and 2-deoxy-D-glucose (50 mM) in C wells. The different glycolysis stress parameters were obtained by subtracting the average respiration rates before and after the addition of the special drugs. The Agilent glycolysis pressure assay report generator (Agilent Technologies, Santa Clara, USA) was used to analyze the pressure changes of cell glycolysis in each experimental group.

#### Pyruvate kinase activity assay

In this experiment, the PKM protein solution (final concentration of 11 nM after dilution) was diluted with Tris-Hcl solution (Sangon Biotech, 1:20 diluted with sterile deionized water at a final concentration of 50 mM, and an appropriate amount of concentrated hydrochloric acid solution was added to keep the PH value at about pH 7.5). The solution was then rapidly vortex for several seconds to fully mix the solution. CLT and Adenosine 5′-triphosphate disodium salt hydrate (Sigma-Aldrich, California, USA) were quickly added to the PKM solution. The final concentration of CLT and Adenosine 5′-triphosphate disodium salt were both 100 μM, and the final concentration of PKM was 10 nM. And the same volume of Tris-HCl solution was added to the Control group. All samples were quickly placed on ice and incubated for 30 minutes. Enzyme activity detection was carried out in strict accordance with the experimental procedure of the Pyruvate kinase activity assay kit (Sigma-Aldrich, California, USA). After 2 to 3 minutes, the initial absorbance measurement was taken at 570 nm (A570)initial. The plate was incubated at 25 ℃ and A570 measurements were taken every 5 minutes. All the 7 measurement data were recorded.

#### ATP**、**NADP+/NADPH**、**GSH and ROS assay

Human A549 cell line & PKM knockout A549 cell line, Human HK-2 cell line & PKM knockout HK-2 cell line were cultured as mentioned above. Changes in ATP, NADPH, ROS and GSH levels in each group were detected using enhanced ATP assay kits (Beyotime, Shanghai, China), NADP+/NADPH assay kits (Beyotime, Shanghai, China), GSH assay kits (Beyotime, Shanghai, China) and ROS assay kits (Beyotime, Shanghai, China) according to the protocol.

#### Western blotting

Tissue and cell extracts were harvested for western blotting as previously described (Zhang et al., 2021). The concentration of protein was quantified by the BCA method (A:B=50:1). The protein was denatured with SDS-PAGE protein loading buffer (metal bath 100°C, 5min). An equivalent amount of denatured protein was loaded to SDS-PAGE gels (10%) and then transferred to PVDF membranes. The membranes were blocked with 5% skimmed milk and washed after the sealing was completed. The membranes were then incubated with PKM rabbit primary antibody (Abcam, 1:2000 dilution) overnight at 4 °C. And beta-actin (Abcam, 1:10000 dilution) was used as the loading control. On the second day, the membranes were washed by PBST (1×) and added with the diluted Goat Anti-Mouse IgG (H+L) HRP (Bioworld TECHNOLOGY, 1:10000 dilution) or Goat Anti-Rabbit IgG (H+L) HRP (Bioworld TECHNOLOGY, 1:10000 dilution) at 37°C for 2 hours. After three times washes, images were acquired using ChemiDoc^TM^ Touch Imaging System (BIO-RAD, California, USA) under dark conditions and the analysis was processed using Image Lab (BIO-RAD, California, USA).

### MULTI OMICS ANALYSIS

#### Bulk RNA sequencing

Mouse model of UUO was conducted as described above. The mice were divided into three groups: the Control group and UUO group was treated with 0.9% NaCl, and UUO + CLT group receiving free CLT (1 mg/kg) was injected intraperitoneally every day. All animals were sacrificed on day 14 and kidneys were harvested for sequencing. The kidney tissue was carefully processed to extract total RNA to ensure high RNA integrity. A total amount of 1 μg RNA per sample was used as input material for the RNA sample preparations. Sequencing libraries were generated using NEBNext® UltraTM RNA Library Prep Kit for Illumina® (NEB, USA) and index codes were added to attribute sequences to each sample. The clustering of the index-coded samples was performed on a cBot cluster generation system using TruSeq PE Cluster Kit v3-cBot-HS (Illumia). After cluster generation, the library preparations were sequenced on an Illumina Novaseq platform and 150 bp paired-end reads were generated. The preprocessed reads were aligned to a reference genome (Mus_musculus_Ensemble_94) using HISAT2 (version 2.0.5, with default parameter), and the gene expression quantification of each group was performed by using the featureCounts software (version 1.5.0-p3, with default parameter).

#### Single cell RNA-sequence (scRNA-seq) data analysis

Digestion of kidney tissue for scRNA-seq was performed by adapting previously reported protocols (Park et al., 2018). Following digestion, the 10x Genomics single-cell 3’ solution was employed for cell sorting and barcode library preparation. The cells were encapsulated in droplets containing barcoded gel beads, enabling cell lysis and reverse transcription of RNA within each droplet. This process yields single-cell cDNA libraries that incorporate both cell and molecular barcodes. Multiplexing of multiple samples was achieved through the addition of sample-specific barcodes during PCR amplification. Following library preparation, the libraries undergo sequencing on a 10x Genomics Chromium single cell gene expression solution platform, adhering to platform-specific protocols. For each sample, the per-cell expression matrix was generated using cellranger count (https://support<u>. 10xgenomics. com/</u>) with mm10 reference. We used Seurat 4.0 R package to perform the data quality control and analysis. More specifically, the cells whose expression count fewer than 200 genes and genes expressing in fewer than 2 cells were removed, while cells were considered apoptotic or injured when mitochondrial genes were > 10 % and also removed. To accurately annotate the cell types, we normalized the read counts of each cell by dividing them by the total read count of that cell, followed by scaling the values by a factor of 10,000 and applying a logarithmic transformation. Subsequently, we performed principal component analysis (PCA) on the normalized expression matrix, using highly variable genes identified through the FindVariableGenes function in Seurat. The top significant principal components were selected for clustering, with a resolution parameter set to 0.5. The resulting cell clusters were visualized using the Uniform Manifold Approximation and Projection (UMAP) method, and the FindAllMarkers function in Seurat was utilized to identify markers specific to each cluster. Finally, we annotated the cell types or subtypes of each cluster based on previously reported cell type markers (Park et al., 2018). Above analysis resulted in the annotation of 13 distinct cell types, including Macrophage, neutrophil, fibroblast, T cell, NK cell, B cell, endothelial cell, collecting duct, podocyte, proximal tubule, ascending loop of Henle, distal convoluted tubule and cortical cell. Cell type percentage analysis, metabolism and reactome analysis were performed by the Cellscope (CapitalBio Technology, Beijing, China)

#### Liquid chromatography-mass spectrometry (LC-MS) proteomics

High-performance, high-throughput liquid chromatography-mass spectrometry (LC-MS) proteomics analysis of mouse tissue was conducted on Orbitrap Exploris 480 (Thermo Fisher Scientific, MA, USA). The tissue samples were homogenized, and proteins were extracted using RIPA buffer (Thermo Fisher Scientific) containing protease inhibitor (Roche). Protein samples were quantified, reduced, digested and desalted as previously reported (Zhang et al., 2024). Proteomic analyses were performed on an Orbitrap Exploris 480 (Thermo Fisher Scientific, MA, USA) with a nano needle integrated chromatography column (OMIC SOLUTION, Shanghai, China). Raw data were processed using Proteome Discoverer software (Thermo Fisher Scientific) against the Mouse Refseq Protein Database containing 21,992 protein entries (UniProt). Search result filters were selected as follows: a t-test was performed to identify proteins with a significantly different abundance (p < 0.05). Proteomics heatmap was analyzed by Proteome Discoverer software (Thermo Fisher Scientific). Protein modification data as well as the CLT binding analysis were analyzed in the Protein modification module of Proteome Discoverer software.

#### STRING analysis of proteomics

Significant altered protein response to CLT treatment in C57BL/6J WT mice and C57BL/6J B6-db/db mice recognized by proteomics analysis was listed in excel and upload to STRING analysis website (https://cn.string-db.org/). The full STRING network indicates active interaction sources including text mining, experiments, databases, co-expression, neighborhood, gene fusion and co-occurrence. The minimum required interaction score was set to medium confidence (0.400). K-means clustering (number of cluster =14) was used to obtain appropriate subclusters.

#### D-glucose-^13^C_6_ isotope labeled targeted metabolomics

Metabolite standard library building was carried out as previously reported (Jang et al., 2018; Liu and Locasale, 2017). Analytical standards (glucose, fumaric acid, isocitric acid, succinic acid, malate, and fructose-6-phosphorytate) were purchased from TargetMol (Boston, USA). Briefly, each standard detection solution was quickly added to the injection bottle and analyzed using LC-MS with Orbitrap Exploris 120 (Thermoscientific, MA, US). The default standard parameter settings were used for both positive ion and negative ion mode detection. Each compound underwent positive and negative scanning to record their secondary mass spectrometry information. MzVault software was used for data collection and processing. And the compound name, molecular formula, mass spectrometry adduct information (such as +H, +NH4, +Na, - H, etc.), charge number, polarity (positive ion mode or negative ion mode) were built for the recognized compound. The newly constructed library was added in Compound Discoverer (Thermoscientific, MA, US) within the following workflow: Stable Isotope Labeling w Metabolika Pathways and ID using Local Databases. The D-Glucose-^13^C_6_ labeled experimental procedure, sample preparation and the sample analysis protocol are the same as described above.

#### D-glucose-^13^C_6_ isotope labeled non-targeted metabolomics

Human A549 cell line & PKM knockout A549 cells were seeded into 6-well plates and five experimental groups were established: Control, CLT, PKM-/-, PKM-/- + CLT (Cell culture medium without glucose was replaced by D-glucose-^13^C_6_ at the beginning of treatment) and unlabeled control group without the D-glucose-^13^C_6_ replacement. CLT (500 nM) was added in the CLT and PKM-/- + CLT for 48 hours. Then cells were harvested for metabolomics analysis with Orbitrap Exploris 120 mass spectrometer (Thermoscientific, MA, US). The samples were run in positive mode with Hypersil GOLD C18 chromatographic column (Thermoscientific, MA, US) and negative mode with Hypersil GOLD HILIC chromatographic column (Thermoscientific, MA, US). Raw data were uploaded to Compound Discoverer (Thermoscientific, MA, US) for compound identification with the following workflow: Stable Isotope Labeling w Metabolika Pathways and ID using Online Databases. The mass spectrum area and related ^13^C exchange percentage for selected compound were analyzed.

#### Statistical analysis

Data are presented as the mean ± standard deviation. The difference among groups was analyzed by Student’s t-test. Statistical analysis was performed using GraphPad Prism software (GraphPad, USA). P values of less than 0.05 were considered to indicate statistically significant differences, and statistical significance was defined as P < 0.05, **P < 0.01, and ***P < 0.001, ****P < 0.0001, with the exact P value marked for each test in the figure.

## Highlights

1. Quantum calculation plus multiple omics identifies PKM as the target of celastrol
2. Celastrol-PKM interaction induces glucose carbon flux U turn and TCA vortex
3. PKM inhibition shows hypoglycemic, anti-obesity and anti-inflammation effect
4. Celastrol-PKM interaction reproduces ATP-PKM interaction in diabetes pathogenesis

## Graphical abstract

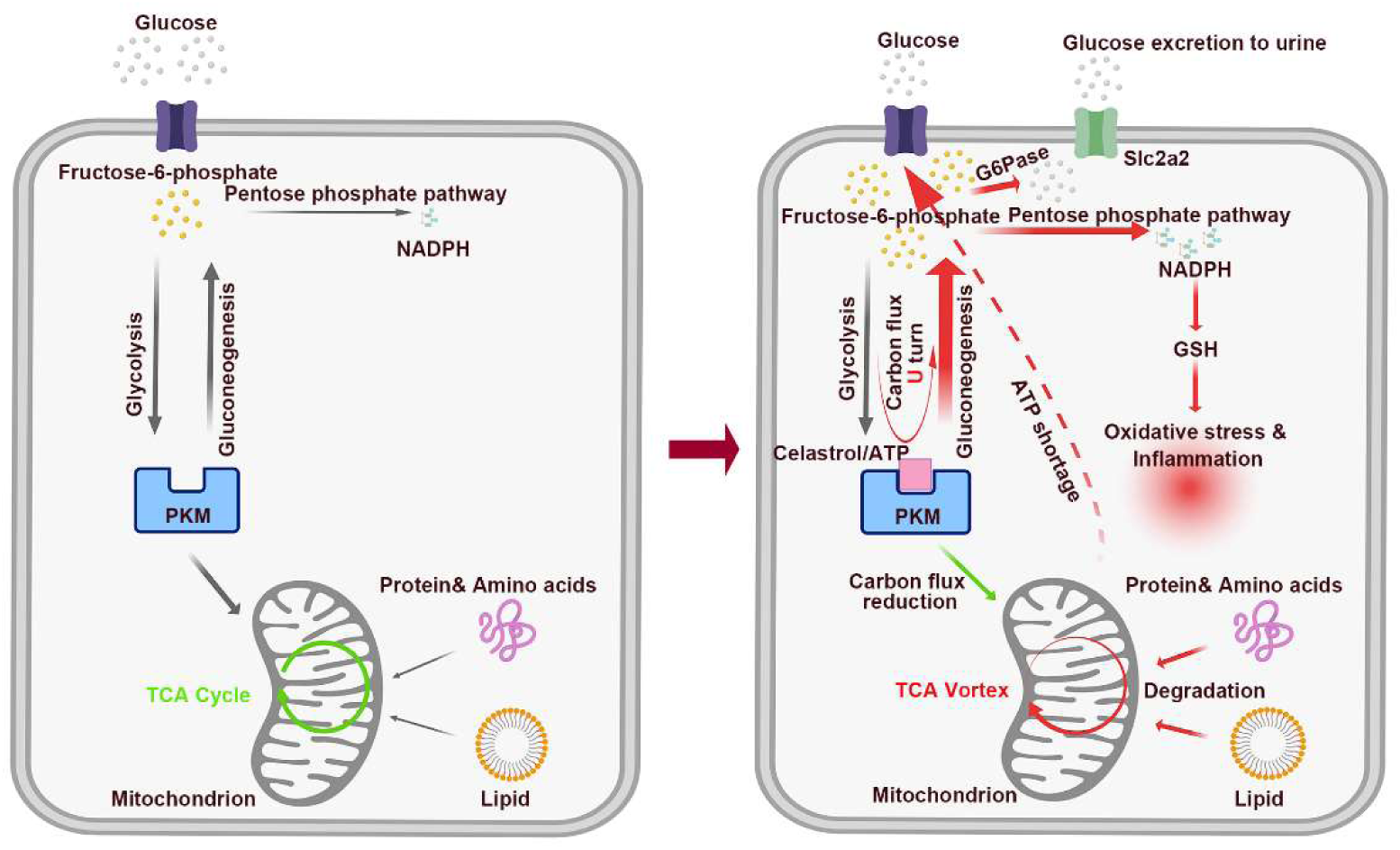

---

**Figure S1.**
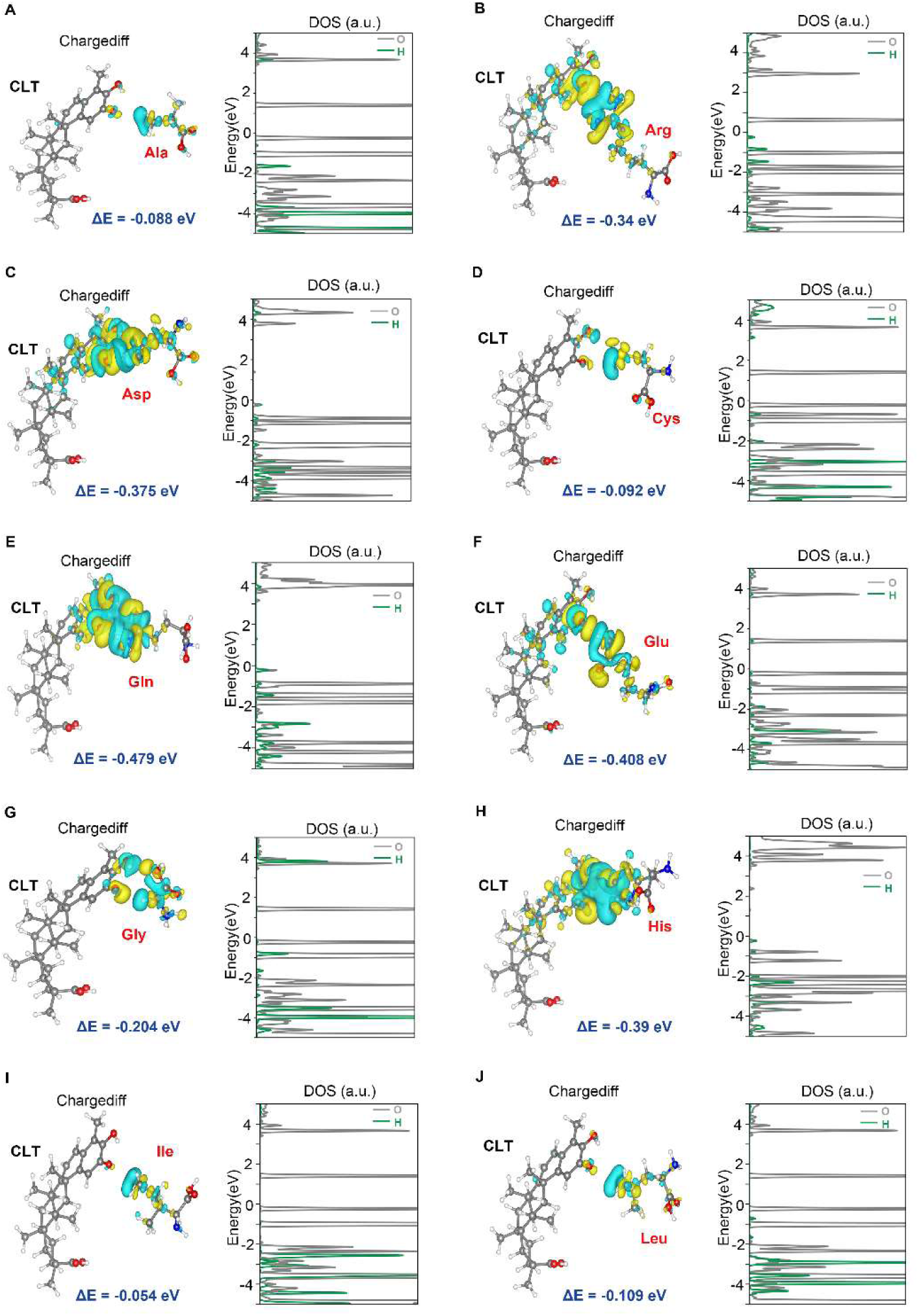
Calculated charge density redistribution, projected density of states and adsorption energy for interaction of selected amino acids and CLT, related to Figure1. The green color denotes electron accumulation, and the yellow color represents depletion. (A) Ala, (B) Arg, (C) Asp, (D) Cys, (E) Gln, (F) Glu, (G) Gly, (H) His, (I) Ile and (J) Leu.

**Figure S2.**
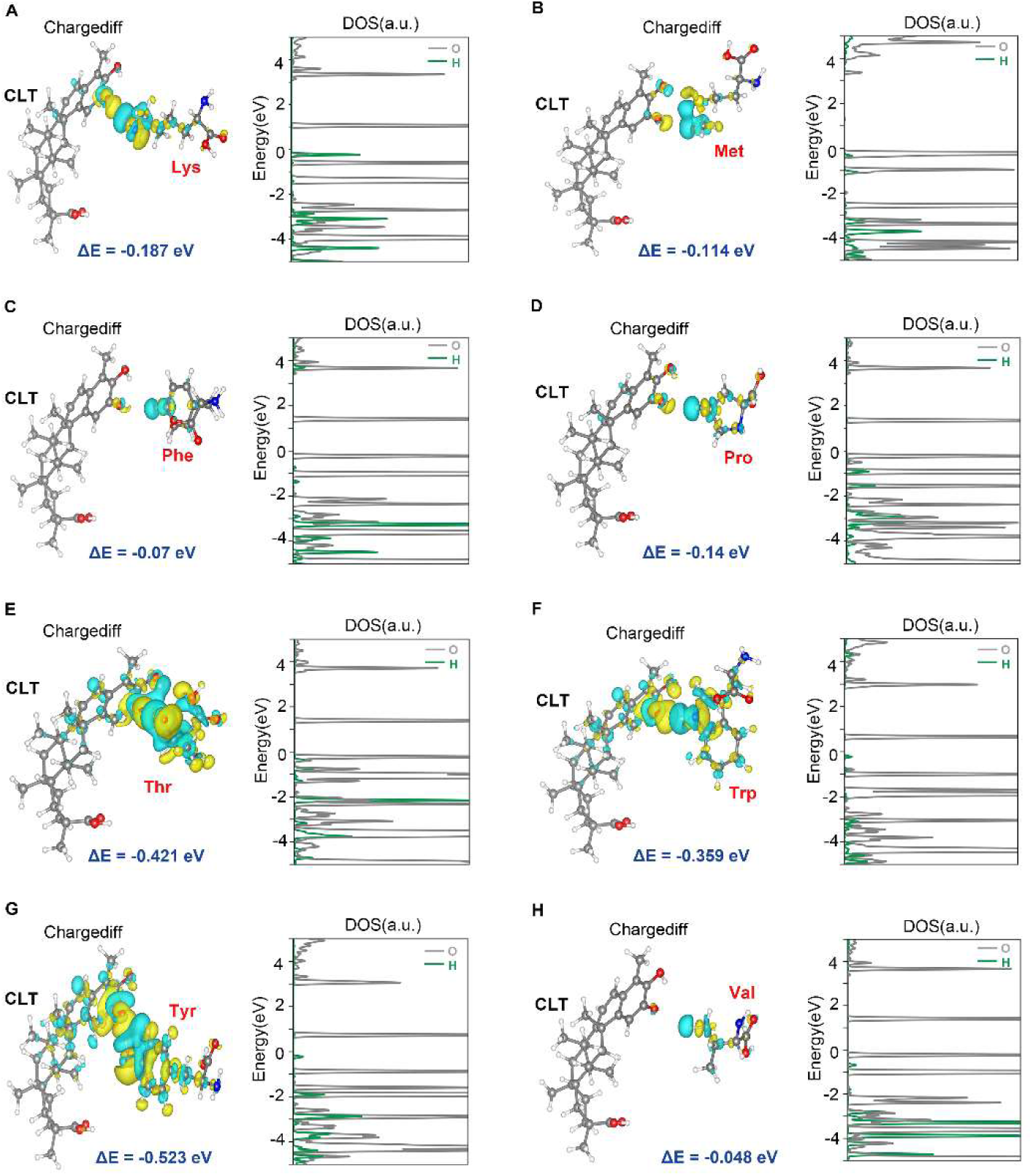
Calculated charge density redistribution, projected density of states and adsorption energy for interaction of selected amino acids and CLT, related to Figure1. The green color denotes electron accumulation, and the yellow color represents depletion. (A) Lys, (B) Met, (C) Phe, (D) Pro, (E) Thr, (F) Trp, (G) Tyr and (H) Val.

**Figure S3.**
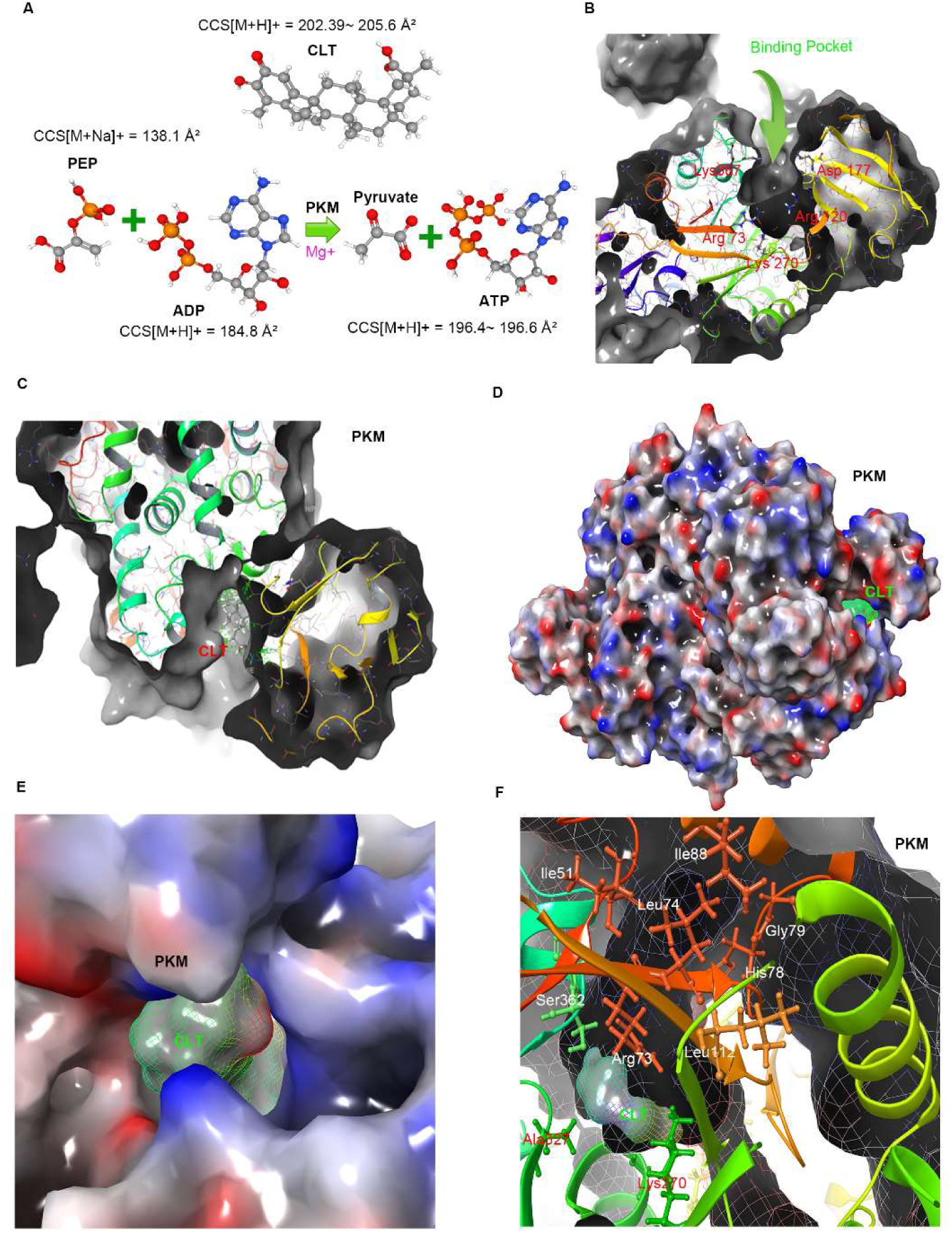
Surface analysis of CLT binding pocket, related to Figure2. (A) The collision cross section of CLT, PEP, ADP & ATP. (B) The surface view of PKM binding pocket without CLT. (C) The surface view of CLT in the binding pocket of PKM. (D) The charge surface of CLT-PKM complex. The blue indicates negative-charged and the red indicates positive-charged on PKM, while the green and the red on CLT denote negative and positive charges, respectively. (E)The charge surface of PKM binding pocket inlet for CLT. The blue indicates negative-charged and the red for positive-charged on PKM, while the green indicates negative-charged and the red for positive-charged on CLT. (F) Amino acid constitution of PKM binding pocket for CLT, including Ile51, Ile88, Leu74, Leu112, Gly79, His78, Ser362 and Arg73.

**Figure S4.**
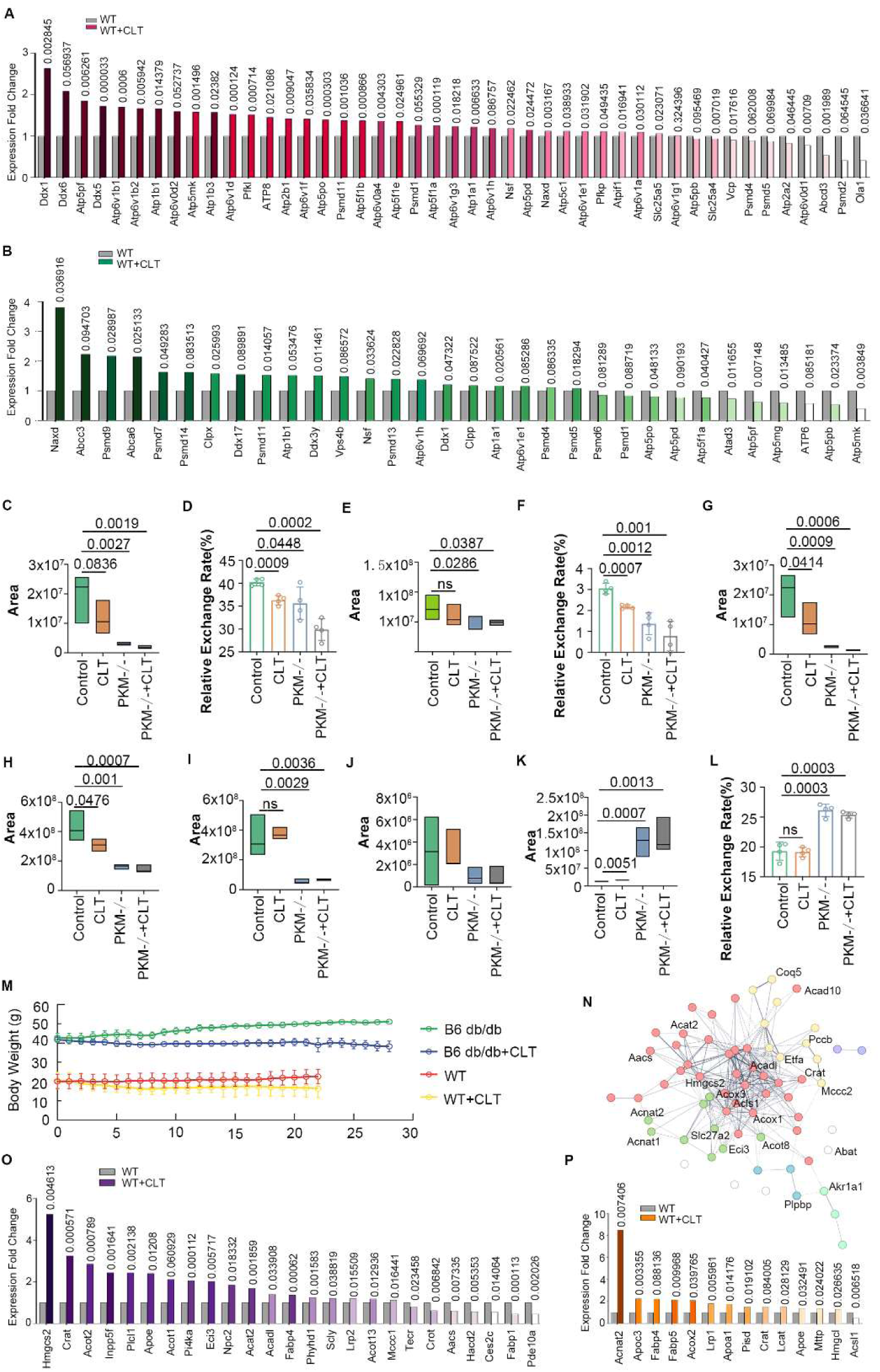
CLT treatment and PKM knockout induce lipid degradation, related to Figure 4, 5 and 6. (A)Expression fold change of proteins related to ATP in 3-week CLT-treated WT mouse kidney. (B)Expression fold change of proteins related to ATP in 3-week CLT-treated WT mouse liver. (C-F) Palmitoylcarnitine, propionylcarnitine and their ^13^C glucose exchange rates were all decreased in CLT, PKM-/- and PKM-/- + CLT group. (G-J) Carnitine, acetyl-L-carnitine, 2-hydroxypropyl stearate, stearate were all decreased in CLT, PKM-/- and PKM-/- + CLT group. (K&L) Glycerophosphocholine and its exchange rate were increased in CLT, PKM-/- and PKM-/- + CLT groups. (M) Body weights decreased in 3-week CLT-treated WT mice (n=5 for WT group, n=6 for CLT group, p<0.0001) and 4-week CLT-treated B6 db/db mice (n=12 for B6-db/db group, n=16 for B6-db/db + CLT group, p<0.0001). (N) Subcluster of total PPI network cluster in Figure 4A related to lipid metabolism. (O) Expression fold change of proteins related to lipid metabolism in 3-week CLT-treated WT mouse kidney. (P) Expression fold change of proteins related to lipid metabolism in 3-week CLT-treated WT mouse liver.

**Figure S5.**
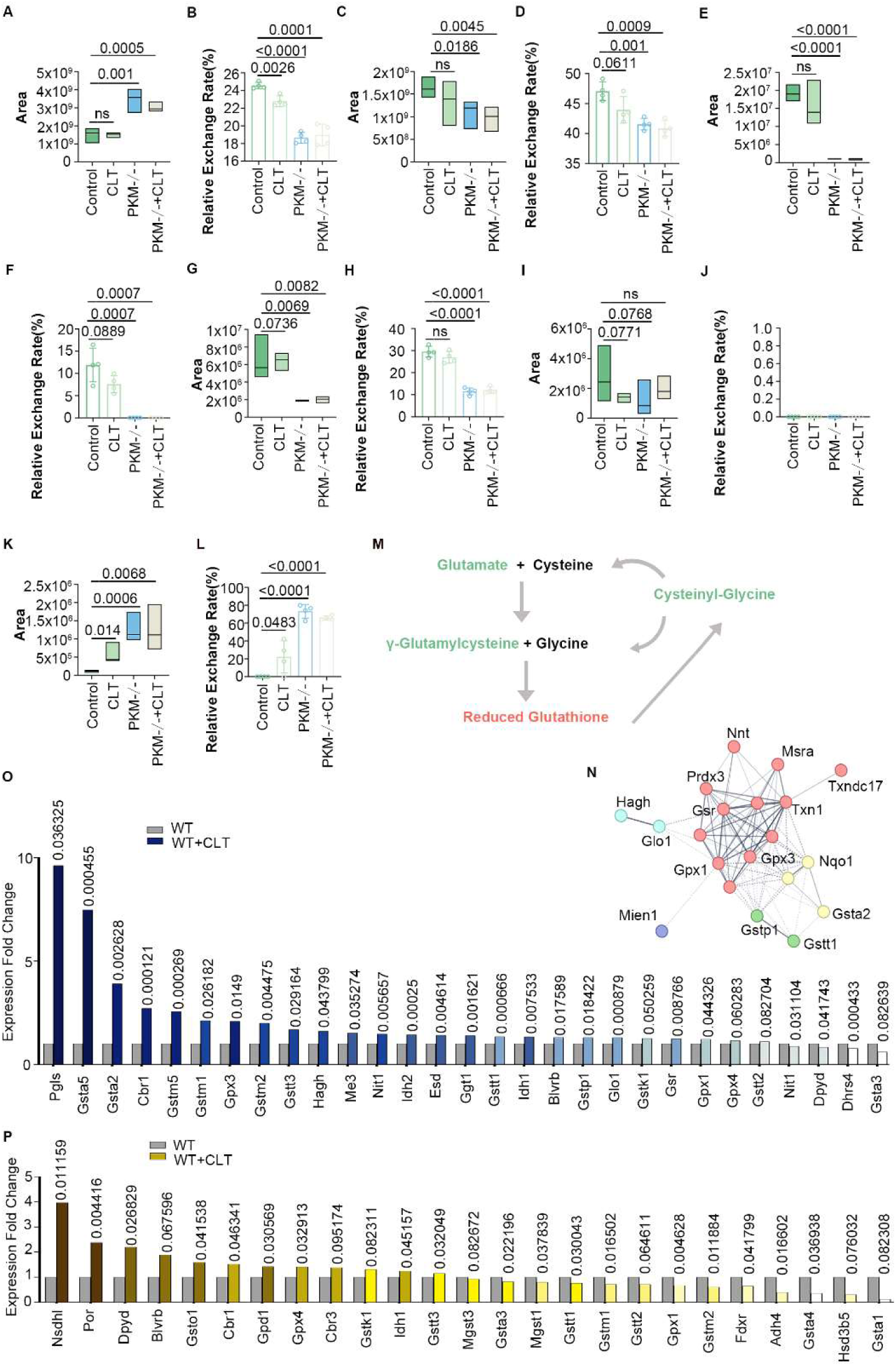
CLT treatment PKM knockout increase the GSH levels by enhancing the PPP, related to Figure 4, 5 and 6. (A&B) GSH levels were increased along with a decreased ^13^C glucose exchange rate in CLT, PKM-/- or PKM-/- + CLT A549 cell. (C-J) Glutamate, glutamine, gamma-glutamylcysteine, cysteinyl-glycine and their ^13^C glucose exchange rate were decreased in CLT, PKM-/- and PKM-/- + CLT A549 cells. (K&L) 6-deoxy-5-ketofructose 1-phosphate and its ^13^C glucose exchange rate were increased in CLT, PKM-/- or PKM-/- + CLT A549 cells. (M) GSH and its ingredient network. The green indicates reduction and the red means upregulation. (N) Subcluster of Figure 4A related to GSH regulation. (O) Proteomics expression fold change related to GSH in 3-week CLT-treated WT mouse kidney. (P) Proteomics expression fold change of proteins related to GSH in 3-week CLT-treated WT mouse liver.

**Figure S6.**
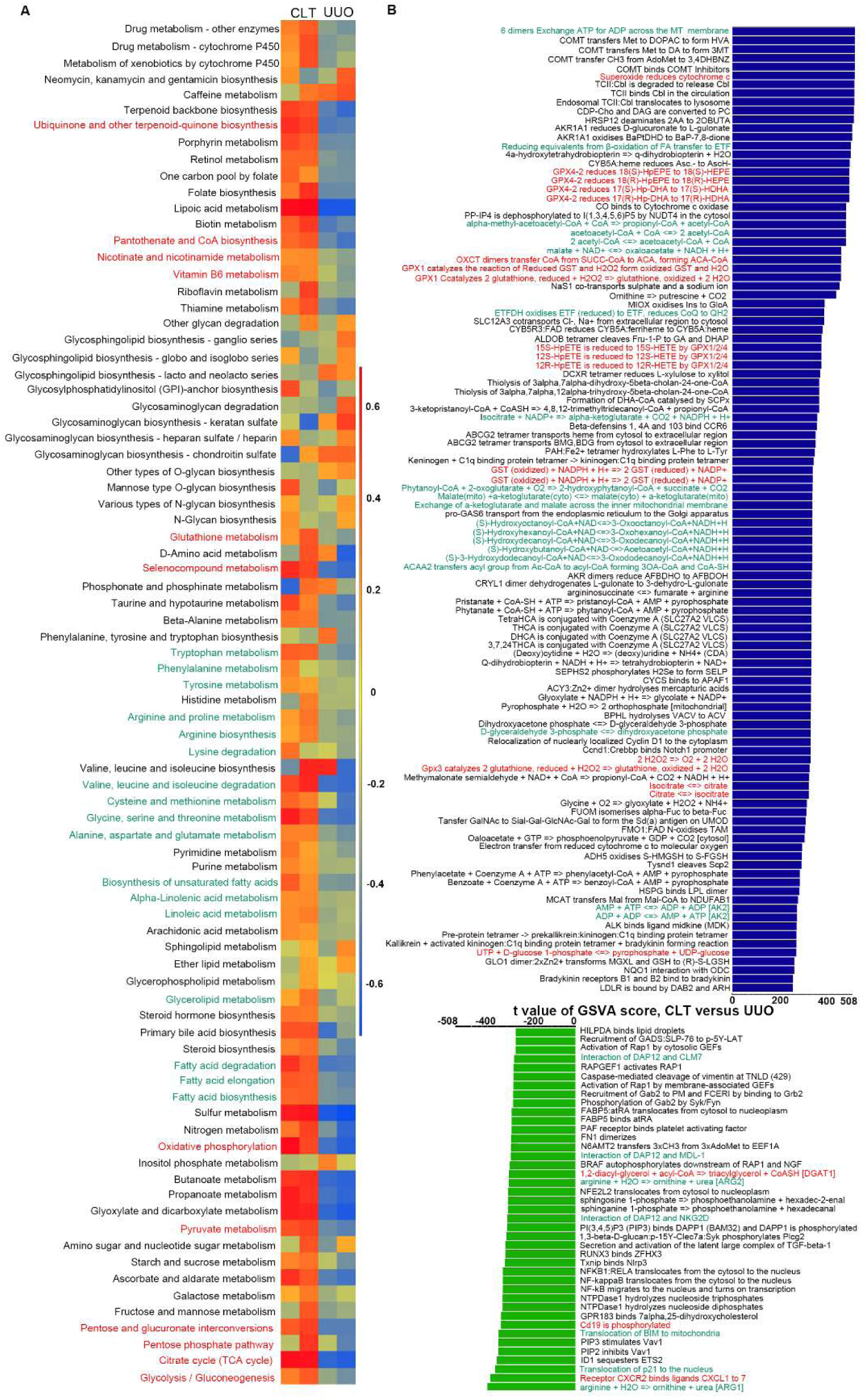
Metabolism and reactome analysis confirmed the biochemical effect of CLT through metabolic and proteomic result, related to Figure 7. (A) Heatmap showing the expression of metabolism-related pathways between CLT and CLT+UUO. (B) Barplot showing the reactome changes after CLT treatment in UUO kidney.

**Figure S7.**
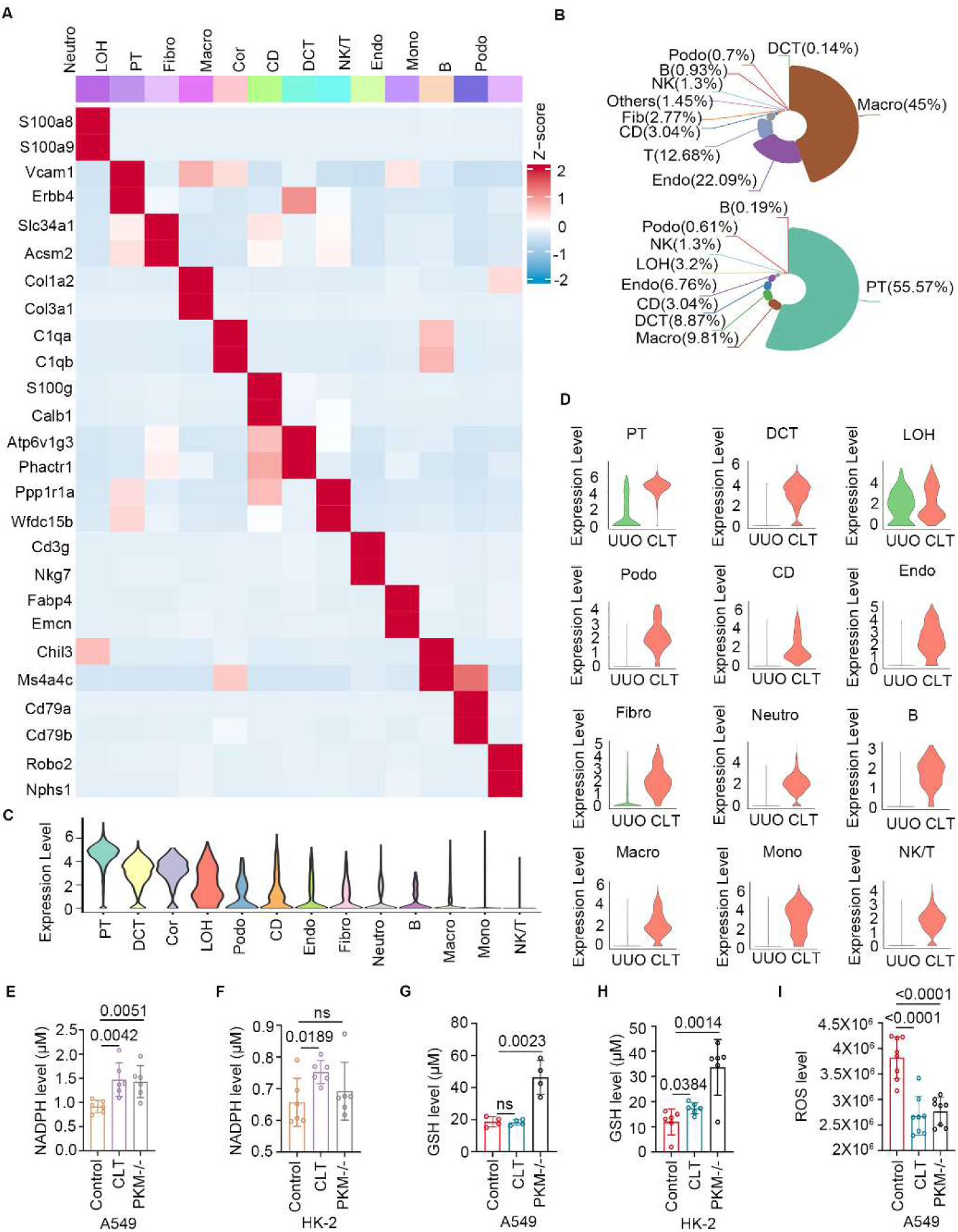
Gpx3 is a key factor of CLT’s anti-inflammatory effect, related to Figure 7. (A) Heatmap showing the gene marker of PT, DCT, LOH, Podo, CD, Endo, Fibro, Neutro, B, Macro, Mono, NK/T cell. (B) Comparison of cell type percentage in UUO kidney and CLT-treated UUO kidney. (C) Violin plot showing the expression of Gpx3 in PT, DCT, LOH, Podo, CD, Endo, Fibro, Neutro, B, Macro, Mono, NK/T cell. (D) The expression of Gpx3 was all increased by CLT treatment in different cell types. (E) CLT and PKM knockout increase NADPH in A549 cells. n=6, p value is listed in the chart. (F) CLT and PKM knockout increase NADPH in HK-2 cells. n=6, p value is listed in the chart. (G) CLT and PKM knockout increase GSH level in A549 cells. n=6, p value is listed in the chart compared to the control group. (H) CLT and PKM knockout increase GSH level in HK-2 cells. n=6, p value is listed in the chart compared to the control group. (I) CLT treatment and PKM knockout reduce the ROS level in A549 cells. n=8, p value is listed in the chart compared to the control group.

